# Interacting Hosts with Microbiome Exchange: An Extension of Metacommunity Theory for Discrete Interactions^*^

**DOI:** 10.1101/2025.07.16.665033

**Authors:** Michael Johnson, Mason A. Porter

**Affiliations:** Department of Mathematics, University of California, Los Angeles, CA 90095, USA; Department of Mathematics and Department of Sociology, University of California, Los Angeles, CA 90095, USA; Santa Fe Institute, Santa Fe, NM 87501, USA

**Author notes:** Submitted to the editors DATE. **Funding:** This work was funded by the National Science Foundation (through award 2124903).

**Keywords:** metacommunity theory, mass effects, microbiomes, mathematical ecology, networks, stochastic differential equations

## Abstract

Microbiomes, which are collections of interacting microbes in an environment, often substantially impact the environmental patches or living hosts that they occupy. In microbiome models, it is important to consider both the local dynamics within an environment and exchanges of microbiomes between environments. One way to incorporate these and other interactions across multiple scales is to employ metacommunity theory. Metacommunity models commonly assume continuous microbiome dispersal between the environments in which local microbiome dynamics occur. Under this assumption, a single parameter between each pair of environments controls the dispersal rate between those environments. This metacommunity framework is well-suited to abiotic environmental patches, but it fails to capture an essential aspect of the microbiomes of living hosts, which generally do not interact continuously with each other. Instead, living hosts interact with each other in discrete time intervals. In this paper, we develop a modeling framework that encodes such discrete interactions and uses two parameters to separately control the interaction frequencies between hosts and the amount of microbiome exchange during each interaction. We derive analytical approximations of models in our framework in three parameter regimes and prove that they are accurate in those regimes. We compare these approximations to numerical simulations for an illustrative model. We demonstrate that both parameters in our modeling framework are necessary to determine microbiome dynamics. Key features of the dynamics, such as microbiome convergence across hosts, depend sensitively on the interplay between interaction frequency and strength.

**MSC codes:** 92-10, 92C42, 92D40, 60J76

## 1. Introduction

The microbiomes of humans and other animals play a critical role in their functioning and health [9, 27, 28], and there is strong evidence that a host’s social interactions significantly impact their microbiome composition [21, 22, 26]. Therefore, it is important to study ecological modeling frameworks that account simultaneously for microbe-scale dynamics and the effects of host interactions [1, 15]. For example, socially determined microbiome signatures are significant indicators of childhood airway development [3], communicable-disease resistance [23], and mental health [10].

### 1.1. Models of Local Ecological Dynamics

The dynamics of microbe populations are affected by environmental factors and the abundances of microbe species. A classical approach to study such populations is to analyze phenomenological dynamical-systems models [6], such as the generalized Lotka–Volterra model [5]

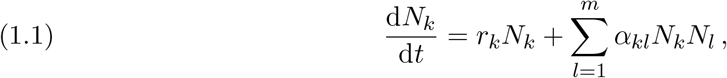

which describes the dynamics of the abundances *N*_1_(*t*), …, *N*_*m*_(*t*) of *m* coexisting microbe species. Each species *k* has an intrinsic birth rate and a death rate, which are combined into a single parameter *r*_*k*_. A positive *r*_*k*_ signifies that the birth rate exceeds the death rate, and a negative *r*_*k*_ signifies that the death rate exceeds the birth rate. Each cross parameter *α*_*kl*_ quantifies the effect of species *l* on the population of species *k*. A positive *α*_*kl*_ indicates that species *l* is beneficial to species *k*, and a negative *α*_*kl*_ indicates that species *l* is harmful to species *k*.

Researchers also employ mechanistic models, such as consumer–resource models [5], to describe microbial population dynamics. One class of such models is niche models

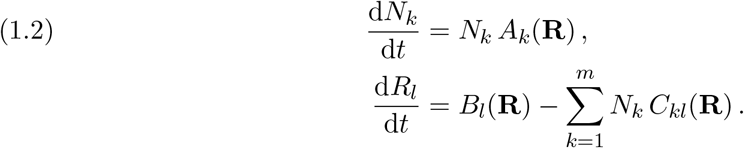

In a niche model, one tracks both the microbe species’ abundances *N*_1_(*t*), *N*_2_(*t*), …, *N*_*m*_(*t*) and the resource abundances *R*_1_(*t*), *R*_2_(*t*), …, *R*_*n*_(*t*). The abundance *N*_*k*_(*t*) of microbe species *k* grows or decays according to a growth rate that is a function *A*_*k*_(**R**) of the resource abundances. The resource abundance *R*_*l*_(*t*) is affected both by the resource abundances and by the consumption of the resource by microbes. The function *B*_*l*_(**R**) encodes the intrinsic dynamics of the resource abundances. The function *C*_*kl*_(**R**) encodes the amount of resource *l* that species *k* consumes per unit abundance of species *k*. Niche models capture the fact that microbes indirectly affect one another through resource competition, rather than interacting directly with each other. Niche models also allow researchers to examine environment-dependent cross-species effects.

### 1.2. Metacommunity Theory

The phenomenological and mechanistic models in subsection 1.1 assume that all microbe species exist in a single environment. Therefore, we refer to these ecological models as models of *local dynamics*. The specification of local ecological dynamics provides a necessary starting point, but it is also necessary to account for interactions across different environments, which are essential to understand microbiome composition in many settings [21, 22, 26]. For example, a coral reef has distinct environmental patches. Stony coral, sponges, algae, and other biotopes provide different conditions to their respective microbiomes. However, the microbiomes of these patches are not isolated and thus impact each other via microbe dispersal [4].

Researchers employ *metacommunity theory* [8, 11] to investigate the effects of multiple-scale interactions that include both local ecological dynamics and interactions across distinct environments. One relevant framework in metacommunity theory is the *mass-effects paradigm* [12, 13, 17, 18, 25], in which researchers study systems with local ecological dynamics and dispersal between environmental patches. Models in this framework are often coupled differential equations of the form

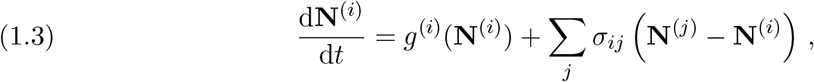

where **N**^(*i*)^(*t*) is a vector that encodes the microbe species abundances in patch *i* at time *t*. For consumer–resource models, one can include resources as separate entries of each vector **N**^(*i*)^(*t*). The autonomous function *g*^(*i*)^ encodes the local dynamics in patch *i*. The parameter *σ*_*ij*_ governs the dispersal between patches *i* and *j*.

### 1.3. Our Contributions

Mass-effects models (e.g., see (1.3)) fail to capture an essential aspect of the microbiomes of many living hosts. Many living hosts (such as humans) do not interact continuously [16] and thus do not sustain a continuous dispersal of microbes. Instead, they interact in discrete time intervals. In the present paper, we develop a framework that considers the discrete nature of host interactions. In this framework, when two hosts interact with each other, they instantaneously exchange some of their microbiomes.

### 1.4. Organization of our Paper

Our paper proceeds as follows. In section 2, we describe our modeling framework for interacting microbiome hosts. In section 3, we describe the behavior of models in our framework in a regime in which hosts interact with low frequency. In section 4, we describe two distinct regimes in which hosts interact with high frequency. In section 5, we present numerical experiments for our framework. In section 6, we discuss conclusions, limitations, and potential future directions of our work. In Appendices A to C, we prove the accuracy of the approximations in sections 3 and 4. Our code is available at https://github.com/mcjcard/Interacting-Hosts-with-Microbe-Exchange.git.

## 2. Our Modeling Framework

In Table 1, we summarize the key notation that we use throughout this paper.

**Table 1.**
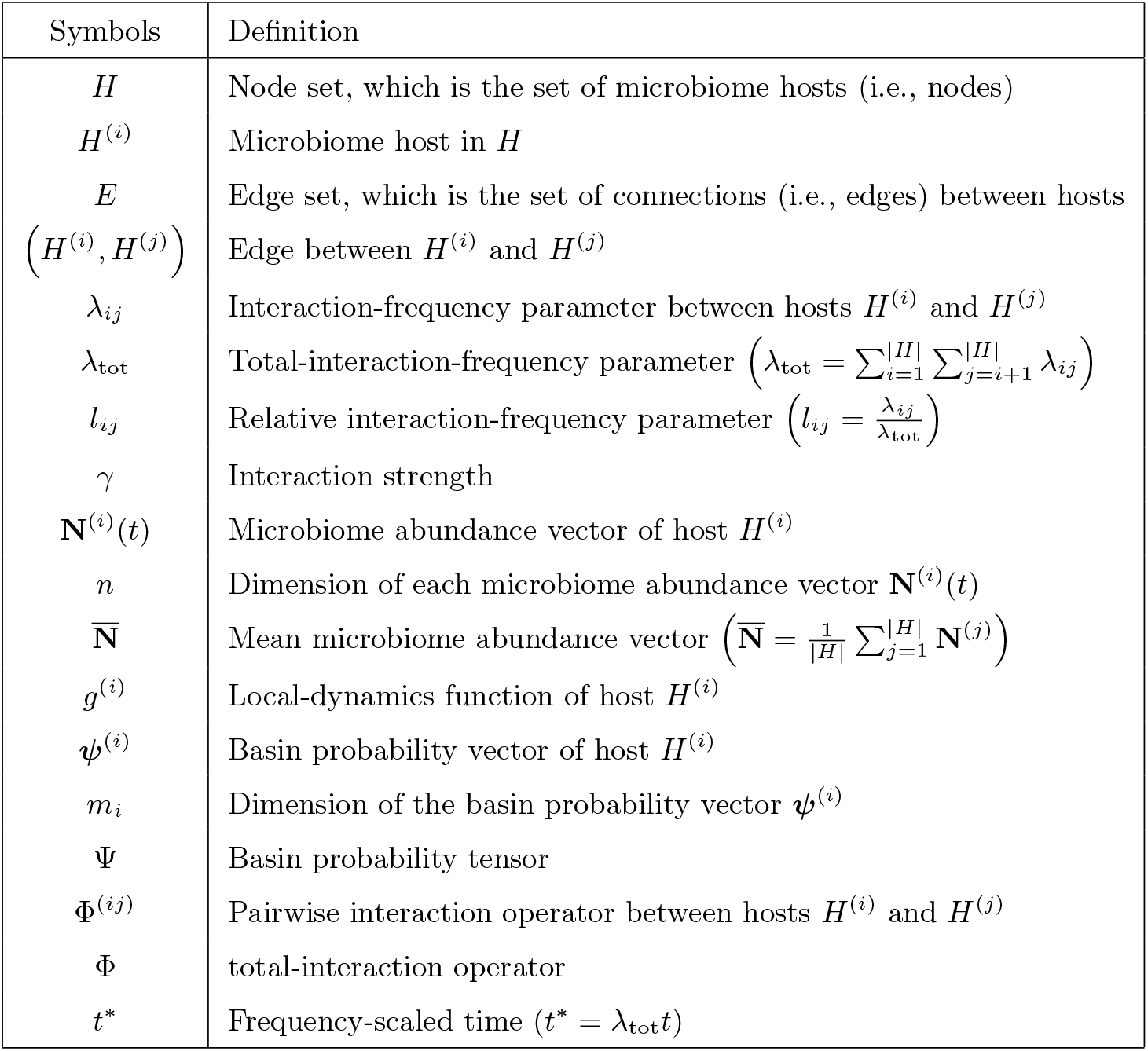
Glossary of our Key Notation.

### 2.1. Interaction Network

To study the microbiomes of living hosts, we consider networks that encode the interactions between these hosts. The simplest type of network is a graph *G* = (*H, E*), which consists of a set *H* of nodes and a set *E ⊆ H × H* of edges between nodes. See [2, 19] for introductions to networks. In the present paper, we use the terms “graph” and “network” interchangeably. Each node of a network represents an entity, and each edge encodes a tie between two entities. In the context of our modeling framework, we refer to each graph as an *interaction network*. The nodes in the node set *H* are microbiome hosts. The edges in the edge set *E ⊆ H × H* encode whether or not two hosts can interact with each other. We represent an undirected edge between two hosts *H*^(*i*)^, *H*^(*j*)^ ∈ *H* by writing either (*H*^(*i*)^, *H*^(*j*)^) or (*H*^(*j*)^, *H*^(*i*)^) (which are equivalent). If an edge exists between two hosts, we say that they are *adjacent*.

Each edge *e* ∈ *E* also has an associated weight *λ*_*e*_ ∈ ℝ^+^. For an edge *e* = (*H*^(*i*)^, *H*^(*j*)^), we equivalently write *λ*_*e*_ or *λ*_*ij*_. If there is no edge between hosts *H*^(*i*)^ and *H*^(*j*)^, we set *λ*_*ij*_ = 0. We refer to this weight as the *interaction-frequency parameter* between hosts *H*^(*i*)^ and *H*^(*j*)^, as it determines the frequency of the interactions between those two hosts. The order of the hosts in indexing is arbitrary, so *λ*_*ji*_ = *λ*_*ij*_. Throughout this paper, any symbol with the index order *ij* is equivalent to that symbol with the reverse index order *ji*. In Figure 1, we show an example of an interaction network with ten hosts. We use this network for our numerical experiments in section 5.

**Figure 1.**
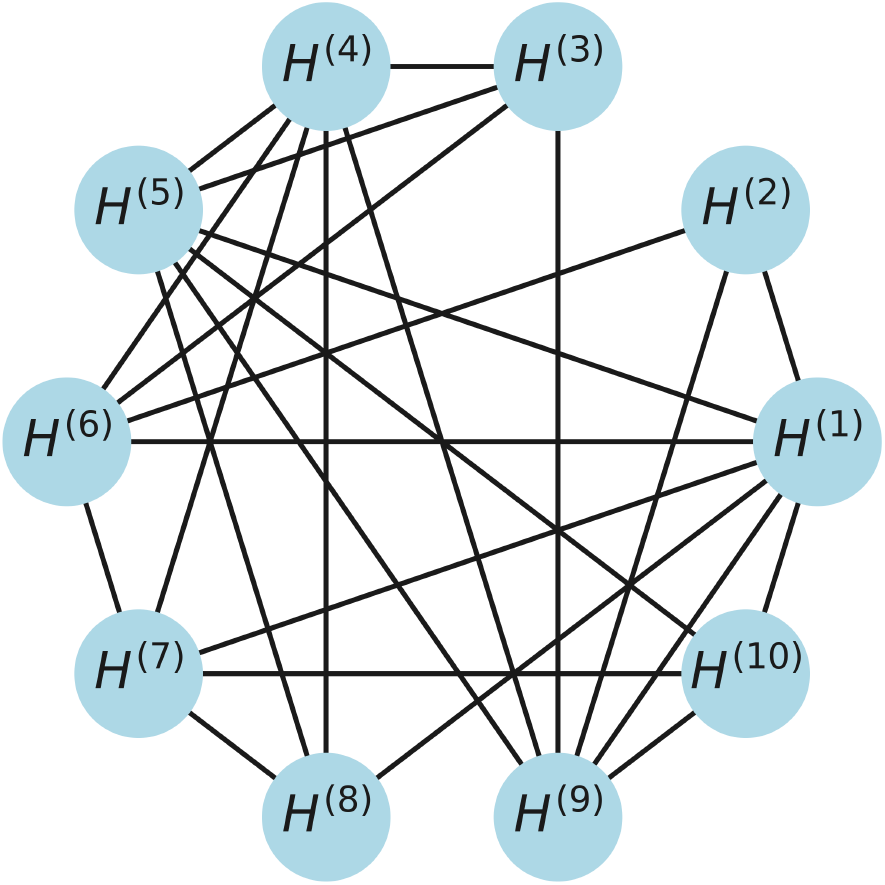
An example of an interaction network with 10 hosts. An edge between two hosts indicates that those two hosts can interact with each other. One can represent heterogeneous interaction-frequency parameters λ_ij_ by using different line widths for different edges. In this example, all λ_ij_ values are either 0 or 1.

### 2.5. Exchange Dynamics

Each host *H*^(*i*)^ ∈ *H* supports a microbiome system. We encode the state of this system with a vector **N**^(*i*)^(*t*) of microbe species abundances. We refer to **N**^(*i*)^(*t*) as the *microbiome abundance vector* of host *H*^(*i*)^. The *k*th entry 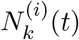 of the microbiome abundance vector encodes the abundance of microbe species *k* in host *H*^(*i*)^ at time *t*. We order the microbiome abundance vector of each host so that its *k*th entry describes the same microbe species for each host. The dimension *n* of all microbiome abundance vectors is the same. If two hosts *H*^(*i*)^ and *H*^(*j*)^ interact at time *t*_*I*_, each host instantaneously exchanges a proportion *γ* of its microbiome with the other host. That is,

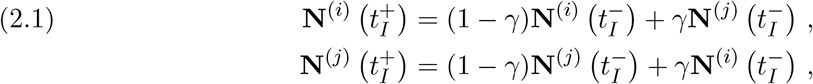

where the parameter *γ* governs the strength of the interaction. At an interaction time *t*_*I*_, the microbiome abundance vector of each host *H*^(*i*)^ satisfies 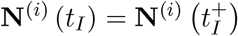. For simplicity, we use the same value of the *interaction strength γ* for each pair of hosts. We model the time between consecutive interactions for a pair of adjacent hosts *H*^(*i*)^ and *H*^(*j*)^ as an exponentially distributed random variable *X*_*ij*_ *∼* Exp(*λ*_*ij*_).

For convenience, we review relevant background information about exponential distributions. For further details, see [7]. The probability density function *f*_*ij*_ for an exponential distribution with parameter *λ*_*ij*_ is

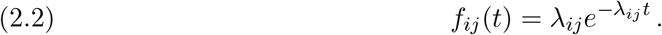

An exponentially distributed random variable *X*_*ij*_ is memoryless. No matter how much time passes after the most recent interaction between hosts *H*^(*i*)^ and *H*^(*j*)^, the time that remains until the next interaction is distributed as *X*_*ij*_. That is,

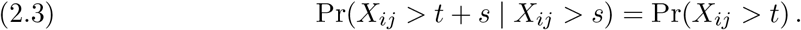

Because *X*_*ij*_ is memoryless, it is easy to describe the random variable (2.4) *X* = min *X*_*ij*_,

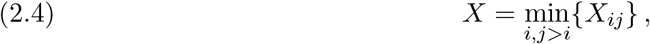

for the time until the next interaction between any pair of hosts. The random variable *X* is also exponentially distributed: *X ∼* Exp(*λ*_tot_), where

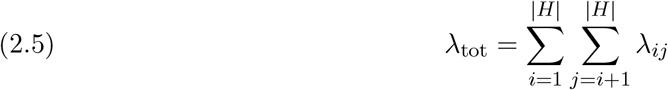

is the *total-interaction-frequency parameter*. The probability that a given interaction is between a specified pair, *H*^(*i*)^ and *H*^(*j*)^, of hosts interacts is the *relative interaction-frequency parameter*

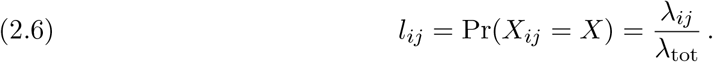

### 2.3. Local Dynamics

Between interactions, an autonomous local dynamical system

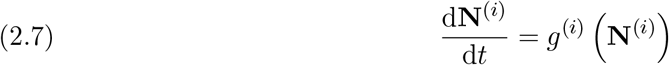

governs the time evolution of the microbiome abundance vector of each host *H*^(*i*)^. This dynamical system encodes the local dynamics of host *H*^(*i*)^. We refer to the function *g*^(*i*)^ as the *local-dynamics function* of host *H*^(*i*)^.

Let the flow **X**^(*i*)^(*t*, **x**) be the solution of

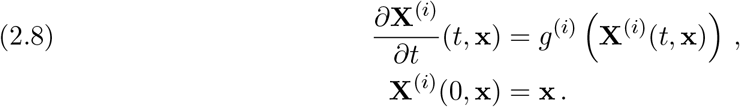

For each *g*^(*i*)^, we require that each element of every valid flow is always finite and nonnegative. Specifically, there is a constant *M* ∈ *R*^+^ such that **x** ∈ [0, *M*]^*n*^ implies that each flow **X**^(*i*)^(*t*, **x**) ∈ [0, *M*]^*n*^ for all times *t ≥* 0. When this condition holds, we say each *g*^(*i*)^ is *bounded*. This is a reasonable assumption for microbiome systems because abundances cannot increase without bound or become negative.

When all *g*^(*i*)^ are bounded, local dynamics cannot cause any microbiome abundance vector **N**^(*i*)^(*t*) to leave the region [0, *M*]^*n*^. Interactions also cannot cause any microbiome abundance vector to leave this region. Consider an interaction at time *t*_*I*_ between hosts *H*^(*i*)^ and *H*^(*j*)^ with microbiome abundance vectors that satisfy 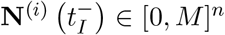 and 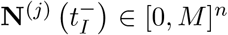. The microbiome exchange between these hosts is an averaging process. After the interaction, the microbiome abundance vectors satisfy 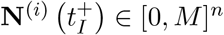 and 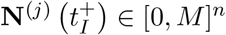. Therefore, if each *g*^(*i*)^ is bounded and each **N**^(*i*)^(0) ∈ [0, *M*]^*n*^, it follows that each **N**^(*i*)^(*t*) ∈ [0, *M*]^*n*^ for all times *t ≥* 0.

We now present an illustrative model of local dynamics that we use repeatedly to illustrate our framework. For this illustrative model, we assume that each host sustains two microbe species. Therefore, each microbiome abundance vector has dimension two. We use the dynamical system

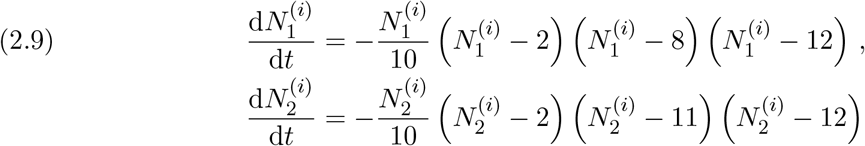

for the local dynamics of each host. In Figure 2, we show this dynamical system’s four stable equilibrium points and their associated basins of attraction. We use the labels 1, 2, 3, and 4 for the basins of attraction of the attractors (2, 2), (12, 2), (2, 12), and (12, 12), respectively.

**Figure 2.**
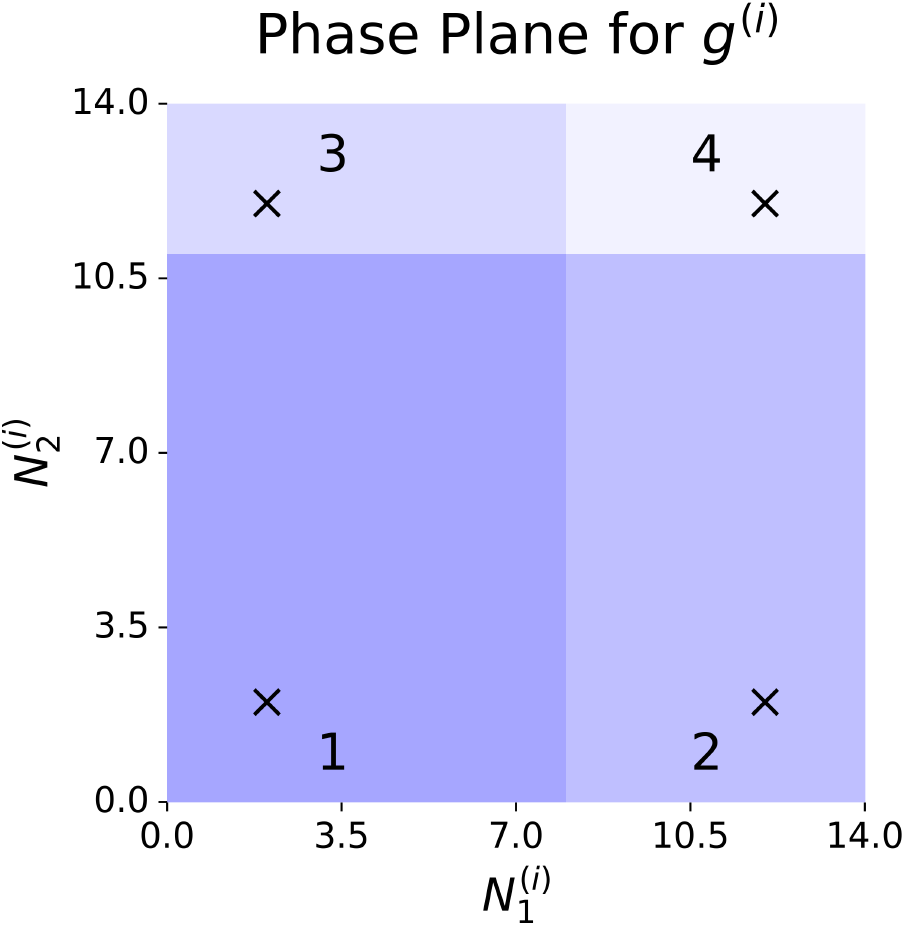
The four stable equilibrium points for the illustrative model (2.9) of local dynamics and their basins of attraction. We use the labels 1, 2, 3, and 4 for the basins of attraction of the attractors (2, 2), (12, 2), (2, 12), and (12, 12), respectively.

Consider an interaction network with two hosts *H*^(1)^ and *H*^(2)^ that are connected by a single edge. Suppose that there is an interaction between the two hosts at time *t*_*I*_ and that 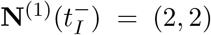 and 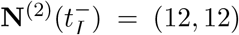. In Figure 3, we show the states of the two hosts immediately after interacting for three values of the interaction strength *γ*. This figure demonstrates that a single interaction can change the basin of attraction of a host’s microbiome abundance vector for sufficiently large *γ*.

**Figure 3.**
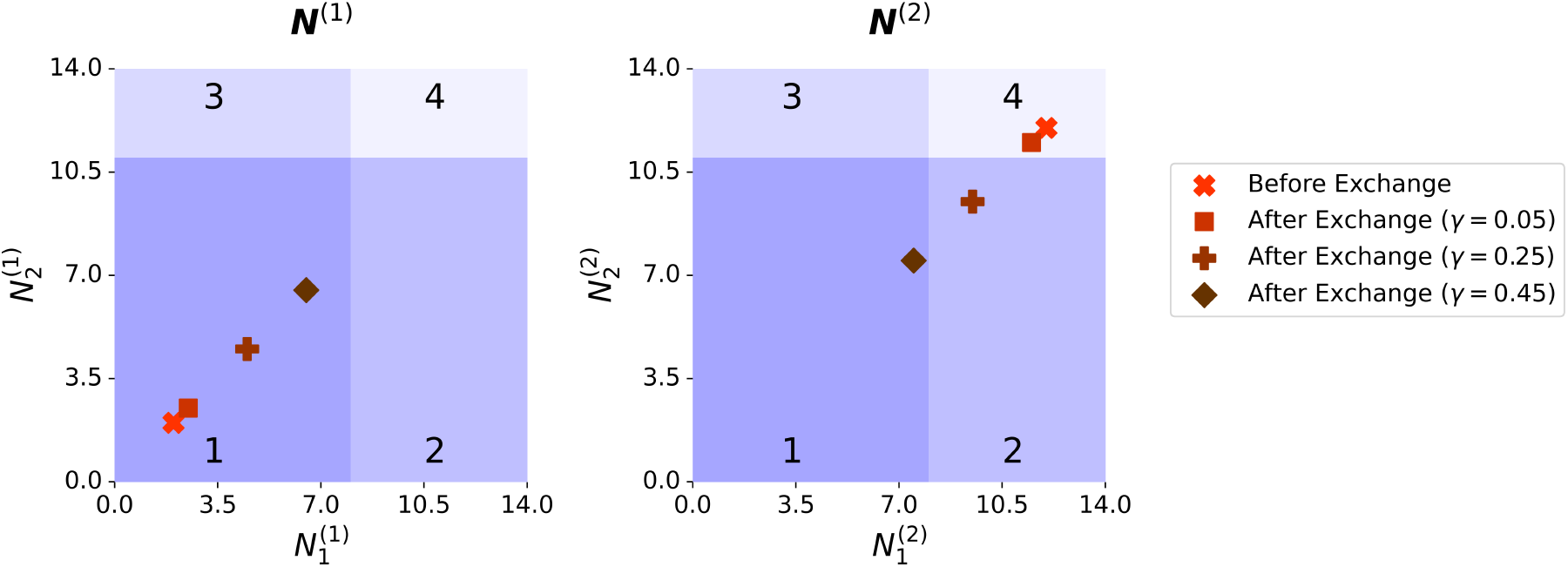
Two hosts with local dynamics (2.9). Immediately before interacting at time t_I_, the hosts have microbiome abundance vectors 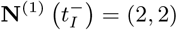 and 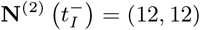. We show the microbiome abundance vectors 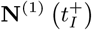 and 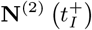 of the two hosts immediately after interacting for interaction strengths γ = 0.05, γ = 0.25, and γ = 0.45.

If interactions occur in sufficiently quick succession, then smaller values of *γ* can also cause transitions in the basin of attraction of a host’s microbiome abundance vector. In Figure 4, we show an example of this phenomenon. We begin with microbiome abundance vectors **N**^(1)^(0) = (2, 2) and **N**^(2)^(0) = (12, 12), and we track the microbiome abundance vectors through five interactions between the hosts. These interactions occur at times 0.1, 0.3, 0.4, 0.7, and 0.73. In this example, the interaction strength is *γ* = 0.32. The first interaction is sufficient to move **N**^(2)^(*t*) from basin 4 to basin 2. However, for **N**^(2)^(*t*) to move from basin 2 to basin 1, two interactions must occur in sufficiently quick succession. In Figure 4, we see that interactions that occur at times 0.3 and 0.4 do not cause this transition. However, interactions at times 0.7 and 0.73 are close enough in time to cause **N**^(2)^(*t*) to move from basin 2 to basin 1.

**Figure 4.**
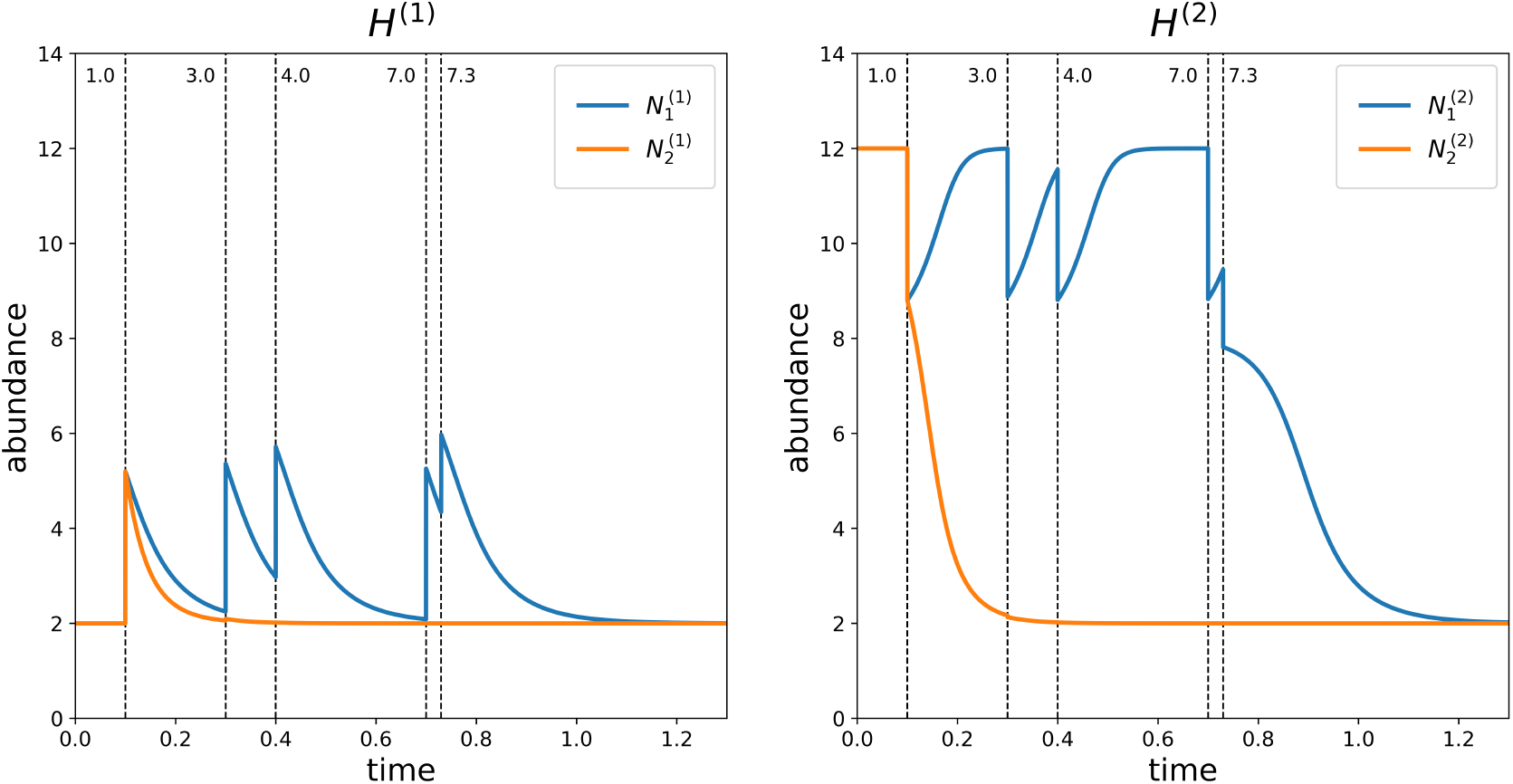
Two hosts with local dynamics (2.9). These hosts have initial states **N**^(1)^(0) = (2, 2) and **N**^(2)^(0) = (12, 12). We show the abundances of microbe species 1 and 2 in each host through the course of five interactions at times 0.1, 0.3, 0.4, 0.7, and 0.73.

## 3. Low-Frequency Approximation (LFA)

In this section, we discuss an approximation that is accurate when all interaction-frequency parameters *λ*_*ij*_ are sufficiently small. We develop this *low-frequency approximation* (LFA) for systems in which the set of stable attractors of each host’s local dynamics is a finite set of equilibrium points. We believe that it is possible to derive extensions of the LFA for systems with other types of attractors, and we discuss this possibility in subsection 6.2. We define relevant terminology in subsections 3.1 and 3.2, and we describe the LFA and outline the proof of its accuracy in subsection 3.3. We give a complete proof in Appendix A.

### 3.1. Basin State Tensor

We illustrated in Figure 3 that interactions can result in transitions of the basin of attraction of a host’s microbiome abundance vector. Local dynamics cannot cause such a transition to occur, so interactions between hosts are necessary for such transitions.

Throughout the rest of section 3, we need to be able to track the basin of attraction of a host’s microbiome abundance vector. To do this, we define a *basin probability vector* ***ψ***^(*i*)^(*t*) for each host *H*^(*i*)^. An entry 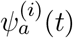 of this vector gives the probability that the microbiome abundance vector of host *H*^(*i*)^ is in basin of attraction *a* at time *t*. If the local dynamics of host *H*^(*i*)^ has *m*_*i*_ basins of attraction, then ***ψ***^(*i*)^(*t*) has dimension *m*_*i*_. Different hosts can have different local dynamics, so the basin probability vectors of different hosts can have different dimensions.

Each local-dynamics function *g*^(*i*)^ is bounded, as described in subsection 2.3. Therefore, for all times *t ≥* 0, each microbiome abundance vector **N**^(*i*)^(*t*) ∈ [0, *M*]^*n*^. Because the set of stable attractors of each host’s local dynamics consists of a finite set of equilibrium points, the set of points 𝒰 ^(*i*)^ *⊂* [0, *M*]^*n*^ that are not in the basin of attraction of some equilibrium point has measure 0. For the LFA to be accurate, we require specific conditions on the local-dynamics functions *g*^(*i*)^ and the interaction strength *γ*. We describe these conditions in the Low-Frequency Approximation Theorem (see Theorem 3.1). When these conditions on *g*^(*i*)^ and *γ* are satisfied and the total-interaction-frequency parameter *λ*_tot_ *→* 0, no microbiome abundance vector **N**^(*i*)^(*t*) lies on the border between basins of attraction at any time *t* in a finite interval [0, *T*] with arbitrarily high probability.

We represent the state of the entire set of hosts using a *basin probability tensor* Ψ(*t*). If each ***ψ***^(*i*)^(*t*) has dimension *m*_*i*_, then Ψ(*t*) has dimension *m*_1_ *×m*_2_ *× … × m* _|*H*|_. An entry of the basin probability tensor 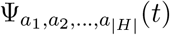 gives the probability that the microbiome abundance vector of each host *H*^(*i*)^ is in basin of attraction *a*_*i*_ at time *t*. If we assume that these probabilities are independent at time *t*, then

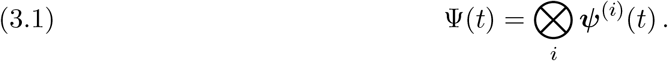

Typically, the ***ψ***^(*i*)^(*t*) are not independent after any interaction, and then (3.1) no longer holds.

### 3.2. Interaction Operators

The basin probability tensor Ψ(*t*) can change only due to an interaction. Unfortunately, knowing only Ψ(*t*) before an interaction and which pair of hosts interacted is insufficient to determine Ψ(*t*) after the interaction. One also needs information about the microbiome abundance vectors of the interacting hosts.

Suppose that an interaction occurs between hosts *H*^(1)^ and *H*^(2)^ at time *t*_*I*_ and that their microbiome abundance vectors 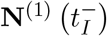 and 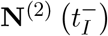 are at stable equilibrium points immediately before the interaction. We make this assumption throughout the rest of this section. If we know that the basins of attraction of 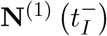 and 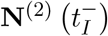 are *a* and *a*, respectively, then we are able to determine the basins of attraction of 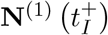 and 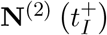. We illustrate this in Figure 5 for an example in which both hosts have local dynamics (2.9) and the interaction strength is *γ* = 0.25. We show 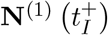 for every possible combination of 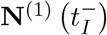 and 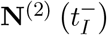. Because 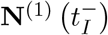 and 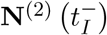 are at stable equilibrium points (by assumption), there are 16 such combinations. For example, for 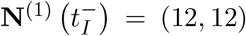, there are four possible values of 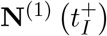. The values are (9.5, 9.5), (12, 9.5), (9.5, 12), and (12, 12); there are four corresponding values ((2, 2), (12, 2), (2, 12), and (12, 12)) of 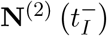. In Figure 5, we mark these four possible values of 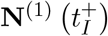 with the diamonds in the upper-right corner. Because host *H*^(2)^ has the same local dynamics as host *H*^(1)^, the situation is identical for 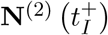, except that we exchange the indices 1 and 2 everywhere.

**Figure 5.**
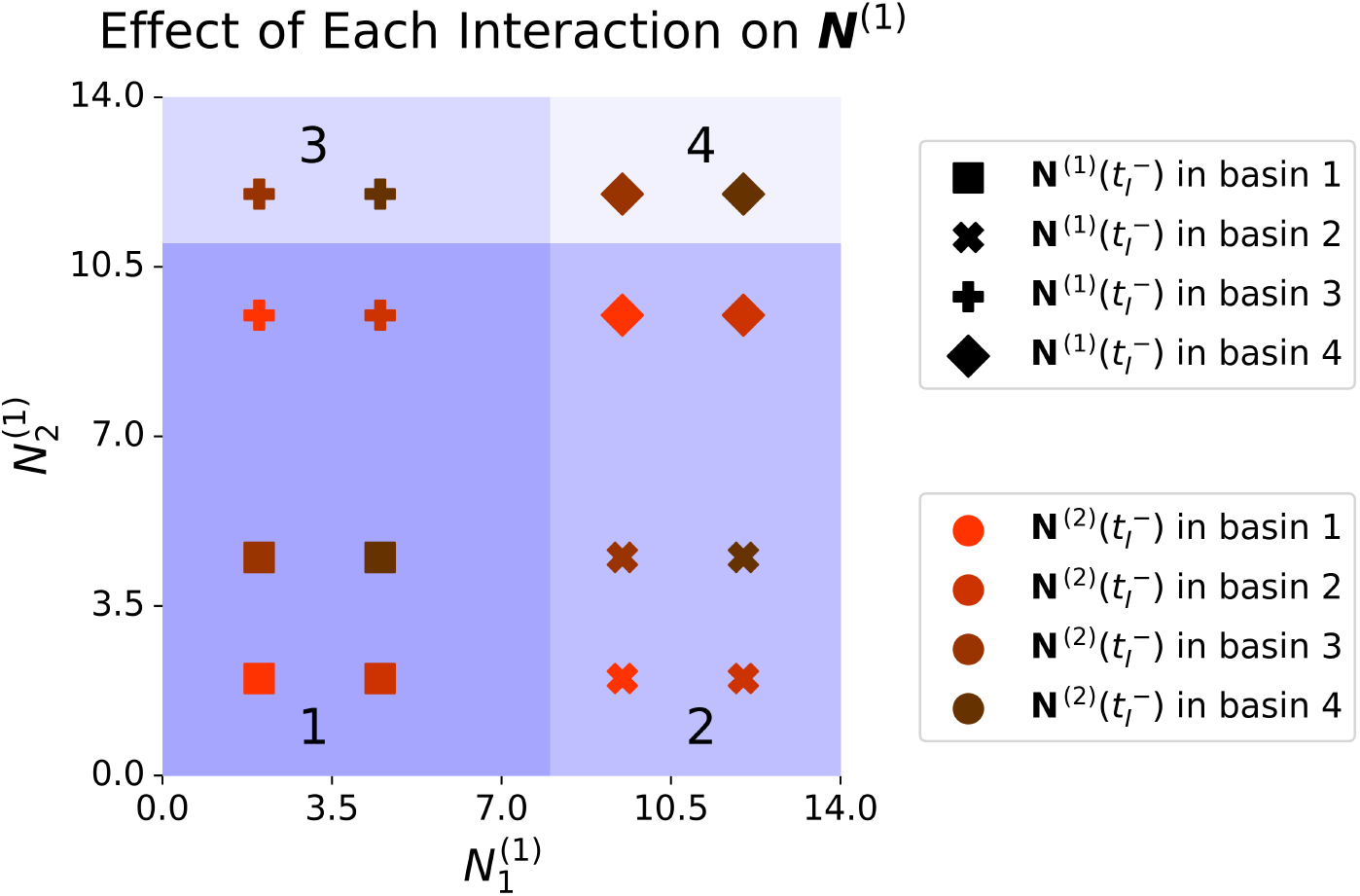
Each possible microbiome abundance vector 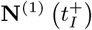 after an interaction at time t_I_ between hosts H^(1)^ and H^(2)^ with local dynamics (2.9), assuming that 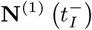 and 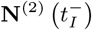 are at stable equilibrium points before the interaction. The marker shapes indicate the basins of attraction of 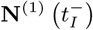, and the marker colors indicate the basins of attraction of 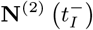. For example, for 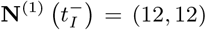, there are four possible values of 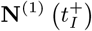. These values are (9.5, 9.5), (12, 9.5), (9.5, 12), and (12, 12); there are four corresponding values ((2, 2), (12, 2), (2, 12), and (12, 12)) of 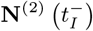. We mark these four possible values of 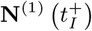 with the diamonds in the upper-right corner. The color indicates the corresponding value of 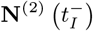.

For some values of the interaction strength *γ*, an interaction between hosts *H*^(*i*)^ and *H*^(*j*)^ can result in either 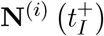 or 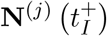 lying on a boundary between basins of attraction, rather than inside a basin of attraction. That is, after such an interaction, we have 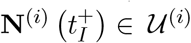 or 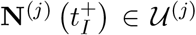. We refer to the set of such interaction strengths as the *boundary set* ℬ_*ij*_ for the hosts *H*^(*i*)^ and *H*^(*j*)^. For the example in Figure 5, the boundary set is ℬ_12_ = {0.1, 0.4}. If hosts *H*^(*i*)^ and *H*^(*j*)^ cannot interact, then ℬ_*ij*_ is the empty set. The *total boundary set* is

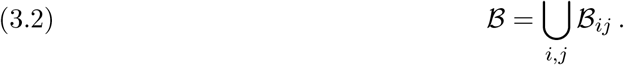

For the LFA, we require *γ* ∉ ℬ, and we assume that this is the case for the rest of this section.

Under our assumptions, we need to know only the basins of attraction of 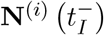 and 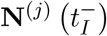 (i.e., before an interaction) to determine the basins of attraction of 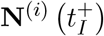 and 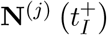 (i.e., after the interaction). Therefore, we need to know only the basin probability tensor 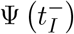 prior to an interaction to determine the basin probability tensor 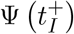 after that interaction. To describe such an interaction-induced change, we define a *pairwise interaction operator* Φ^(*ij*)^ with dimension *m*_1_ *× … × m*_|*H*|_ *× m*_1_ *× … × m*_|*H*|_. Its entry 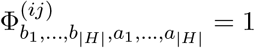 if *b*_*i*_ and *b*_*j*_ are the basins of attraction of 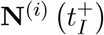 and 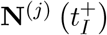, respectively, when *a*_*i*_ and *a*_*j*_ are the respective basins of attraction of 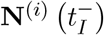 and 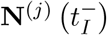 and *b*_*k*_ = *a*_*k*_ for all *k*∉ {*i, j*}. Otherwise, 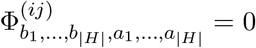.

Consider a two-host interaction network in which both hosts have local dynamics (2.9) and the interaction strength is *γ* = 0.25. The basin probability tensor of this system has dimension 4 *×* 4. Therefore, the pairwise interaction operator Φ^(12)^ has dimension 4 *×* 4 *×* 4 *×* 4. In Figure 5, we show the results of all possible interactions between these two hosts. For example, if 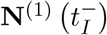 and 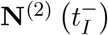 are in basins of attraction 1 and 4, respectively, then the resulting basins of attraction of 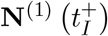 and 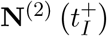 are 1 and 2, respectively. Therefore, 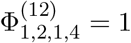. Because there are 16 possible combinations of 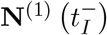 and 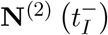, there are 16 entries of Φ^(12)^ that equal 1. We list these 16 entries in Figure 6.

**Figure 6.**
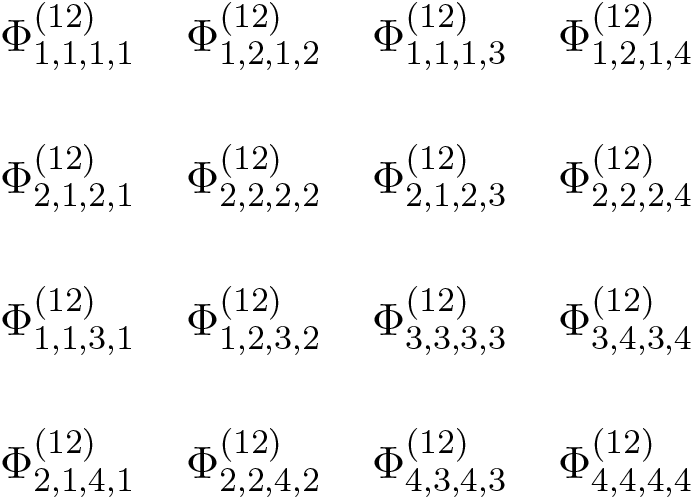
The 16 entries of the pairwise interaction operator that equal 1 for a two-host interaction network in which both hosts have local dynamics (2.9) and the interaction strength is γ = 0.25.

We use the Einstein summation convention [24] to describe how Φ^(*ij*)^ operates on the basin probability tensor. In this convention, one sums over any repeated index that occurs in a single term. For example, we describe the product

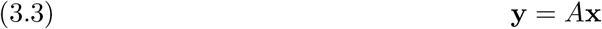

of a matrix *A* and a vector **x** by writing

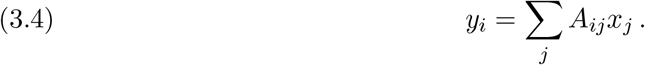

Using the Einstein summation convention, we write

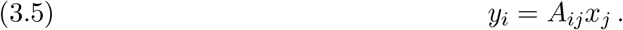

We write the effect of the pairwise interaction operator Φ^(*ij*)^ on the basin probability tensor Ψ as

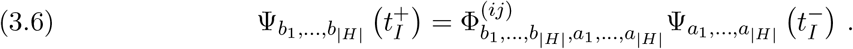

Each Φ^(*ij*)^ is a linear operator on the basin probability tensor. At any time, the next interaction is between hosts *H*^(*i*)^ and *H*^(*j*)^ with probability *l*_*ij*_ (see (2.6)). This yields the *total-interaction operator*

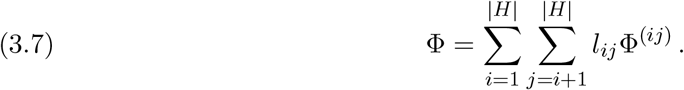

If we now assume that an interaction occurs between some pair of hosts at time *t*_*I*_ (with probabilities *l*_*ij*_ for each pair) and that all 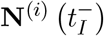 are at a stable equilibrium point, then we obtain

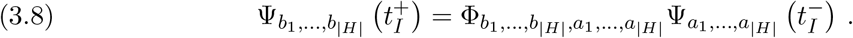

### 3.3. Low-Frequency-Approximation Theorem

The LFA encodes the evolution of the system (2.1, 2.7) when the local dynamics of each host is much faster than the exchange dynamics between hosts. In this regime, each microbiome abundance vector **N**^(*i*)^(*t*) becomes close to a stable equilibrium point before the next interaction that involves host *H*^(*i*)^. Therefore, the total-interaction operator Φ accurately describes the dynamics of the basin probability tensor Ψ(*t*).

Before we state the LFA Theorem, we introduce some helpful terminology. The expected number of host interactions in a time interval of duration Δ*t* is *λ*_tot_Δ*t*. We refer to *t*^*\**^ = *λ*_tot_*t* as the *frequency-scaled time*. We say that the local-dynamics function *g*^(*i*)^ is *inward pointing* at a point **x** if there exists a constant *δ >* 0 such that ∥**y** − **x**∥_2_ *≤ δ* implies that *g*^(*i*)^(**y**)·(**x**−**y**) *>* 0.

#### Theorem 3.1

(Low-Frequency-Approximation Theorem). *Suppose that the attractors of each host’s local dynamics consist of a finite set of stable equilibrium points at which the local-dynamics function g*^(*i*)^ *is inward pointing, and let each g*^(*i*)^ *be continuous and bounded (see* subsection 2.3*). Fix γ*∉ ℬ, *all l*_*ij*_, *and a frequency-scaled time T*^*\**^. *As λ*_tot_ *→* 0, *the basin probability tensor* Ψ(*t*^*\**^) *converges uniformly to* 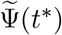 *on* [0, *T*^*\**^], *where*

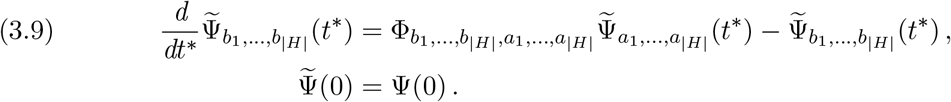

We provide key steps of the proof of Theorem 3.1 in this section. We give a full proof in Appendix A.

As we described in subsection 3.2, if each 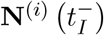 is at a stable equilibrium point, then the interaction operator Φ describes the update of the basin probability tensor after an interaction at time *t*_*I*_. We can construct neighborhoods around each stable equilibrium point of each host’s local dynamics such that if each 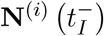 is in one of these neighborhoods, then Φ perfectly describes the transition of Ψ(*t*) due to an interaction at time *t*_*I*_. In Figure 7, we show an example of what these neighborhoods can look like around the stable equilibrium points for our illustrative model (2.9) of local dynamics.

**Figure 7.**
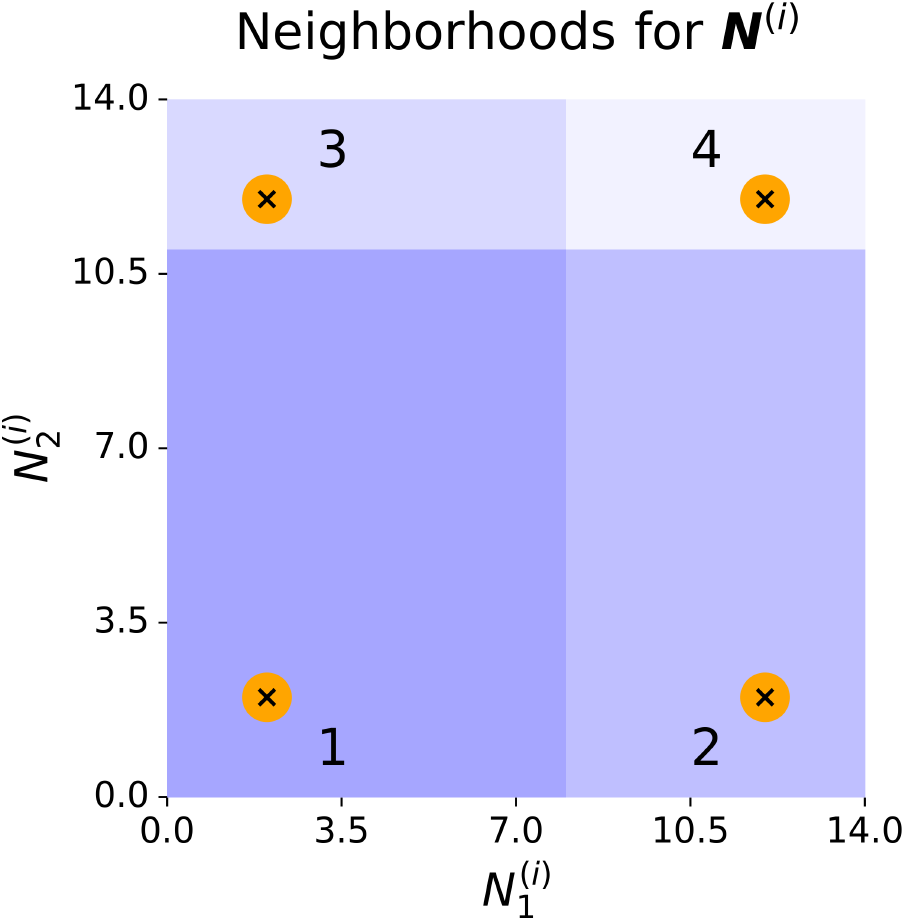
An illustration of potential neighborhoods around the four stable equilibrium points for a host with the local dynamics (2.9). These neighborhoods illustrate potential neighborhoods from Theorem 3.1; they are not the neighborhoods for any particular value of the interaction strength γ.

After an interaction, there is an upper bound on the time that it takes for each microbiome abundance vector to re-enter a neighborhood around a stable equilibrium point. Because the system is finite, there is an upper bound *τ* such that if no interactions occur in the time interval [*t, t* + *τ*], each microbiome abundance vector **N**^(*i*)^(*t* + *τ*) is in one of these neighborhoods. If no two interactions occur within time *τ* of each other, then successive applications of the total-interaction operator Φ perfectly describe the evolution of the basin probability tensor Ψ(*t*). In the frequency-scaled time interval [0, *T*^*\**^], the expected number of system interactions is *T*^*\**^. As *λ*_tot_ *→* 0, the frequency-scaled time *τ*^*\**^ *→* 0. Therefore, it becomes vanishingly unlikely that any pair of interactions occurs within a frequency-scaled time that is less than *τ*^*\**^. Consequently, as *λ*_tot_ *→* 0, the effect on the basin probability tensor Ψ(*t*^*\**^) of all interactions in [0, *T*^*\**^] is described perfectly by the total-interaction operator Φ with arbitrarily high probability. Because the interactions are exponentially distributed, the basin probability tensor Ψ(*t*^*\**^) converges uniformly to 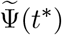 (see (3.9)).

## 4. High-Frequency Approximations

In this section, we discuss two approximations that are accurate for different regimes with large *λ*_tot_. The first of these approximations is the *high-frequency, low-strength approximation* (HFLSA). The HFLSA becomes increasingly accurate as *λ*_tot_ *→ ∞* and *γ →* 0 for fixed relative interaction-frequency parameters *l*_*ij*_ and fixed *λ*_tot_*γ*. This approximation results in a model that has the same form as the mass-effects model (1.3). The second approximation is the *high-frequency, constant-strength approximation* (HFCSA). This approximation becomes increasingly accurate as *λ*_tot_ *→ ∞* for fixed relative interaction-frequency parameters *l*_*ij*_ and fixed interaction strength *γ*.

### 4.1. High-Frequency, Low-Strength Approximation (HFLSA)

The HFLSA is accurate when the interactions between hosts are very frequent but very weak. In this regime, the expectation of the exchange dynamics (2.1) is constant, but the variance of the exchange dynamics is small. This results in an approximate model for the dynamics of each microbiome abundance vector **N**^(*i*)^(*t*) that is deterministic and has terms that encode the effects of the local dynamics and the exchange dynamics. This approximate model has the same form as the mass-effects model (1.3).

#### Theorem 4.1

(High-Frequency, Low-Strength Approximation Theorem). *Fix the relative interaction-frequency parameters l*_*ij*_, *the product λ*_tot_*γ, and a time T. Let each local-dynamics function g*^(*i*)^ *be continuously differentiable and bounded (see subsection* 2.3*), and let ε* ∈ (0, 1] *and δ >* 0 *be arbitrary but fixed. For sufficiently large λ*_tot_, *each host’s microbiome abundance vector* **N**^(*i*)^(*t*) *satisfies*

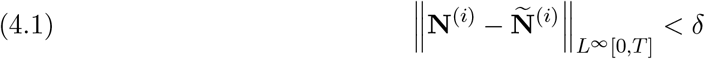

*with probability larger than* 1 − *ε, where*

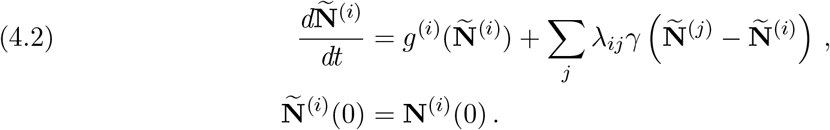

We provide key steps of the proof of Theorem 4.1 in this section. We give a full proof in Appendix B.

Consider the evolution of a microbiome abundance vector **N**^(*i*)^(*t*) over a short time interval [*t*^*′*^, *t*^*′*^ + *dt*]. Without interactions, the effect of the local dynamics over this interval is

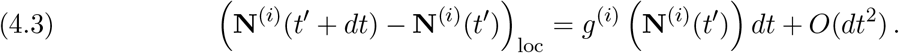

Let

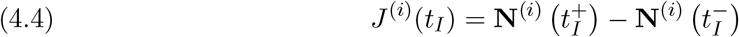

be the effect of an interaction at time *t*_*I*_ on **N**^(*i*)^(*t*). If the interaction does not involve *H*^(*i*)^, then *J*^(*i*)^(*t*_*I*_) = **0**. Otherwise, if the interaction is between *H*^(*i*)^ and *H*^(*j*)^, then

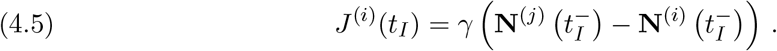

Suppose that no interactions occur precisely at times *t*^*′*^ or *t*^*′*^ + *dt* and that *L* interactions occur during the interval (*t*^*′*^, *t*^*′*^ + *dt*). We denote this ordered set of interactions by 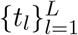. Suppose for all *i* that the microbiome abundance vector **N**^(*i*)^(*t*) changes very little over the interval [*t*^*′*^, *t*^*′*^ + *dt*] such that each *J*^(*i*)^(*t*_*l*_) is well-approximated by

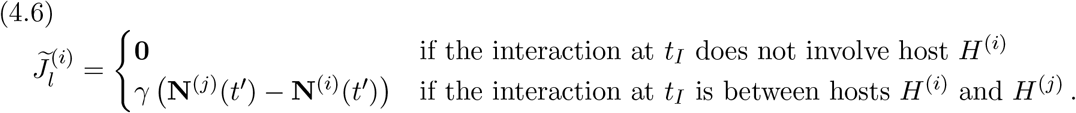

It then follows that the effect of the interactions on the microbiome abundance vector **N**^(*i*)^(*t*) during this interval is well-approximated by

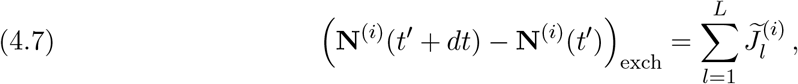

which we henceforth call the *approximate interaction effect*.

Each of the approximate interaction effects 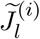 is vector-valued. We denote entry *x* of this vector by 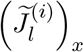. The sum (4.7) is also vector-valued, and we denote entry *x* of this sum by 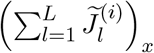. We now calculate the expectation and the variance of each entry of (4.7). The stochasticity in (4.7) arises both from the number *L* of interactions and from which pair of hosts interact at each time. The number of interactions follows a Poisson distribution with mean *λ*_tot_*dt*. The approximate interaction effects 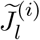 are independent of one another. An interaction at time *t*_*l*_ is between hosts *H*^(*i*)^ and *H*^(*j*)^ with probability *λ*_*ij*_*/λ*_tot_. Therefore,

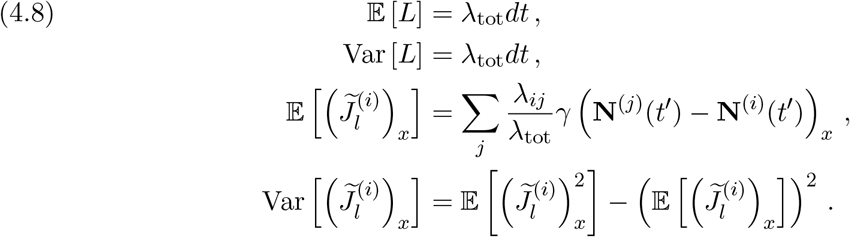

The expectation of each entry of the sum is

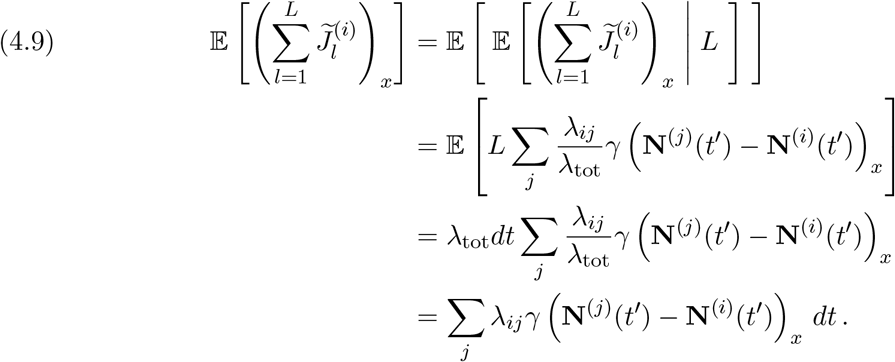

Applying the law of total variance, the variance of each entry of the sum is

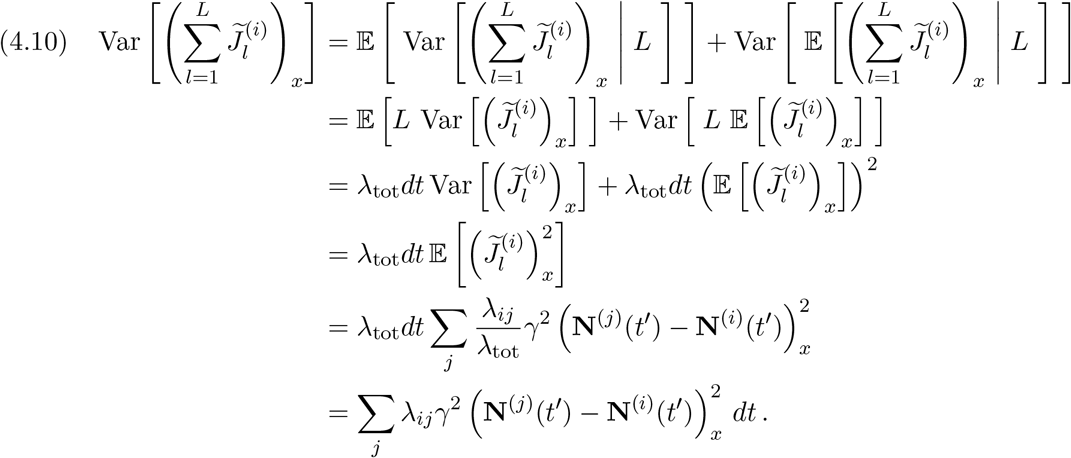

Because *λ*_tot_*γ* is fixed, the interaction strength *γ →* 0 as *λ*_tot_ *→ ∞*. As *γ →* 0, the expectation (4.9) of each entry of (4.7) remains fixed, but the variance (4.10) of each entry decreases to 0. For sufficiently small *γ*, the effect of interactions on the microbiome abundance vector **N**^(*i*)^(*t*) over the interval [*t*^*′*^, *t*^*′*^ + *dt*] is

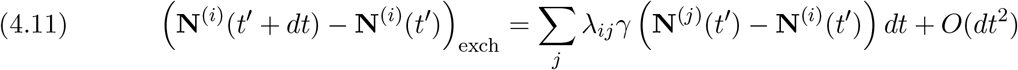

with arbitrarily high probability.

For sufficiently small *γ*, the change in the microbiome abundance vector **N**^(*i*)^(*t*) over the interval [*t*^*′*^, *t*^*′*^ + *dt*] is approximately equal to the sum of the effect (4.3) of local dynamics and the effect (4.11) of interactions. In Appendix B, we show that

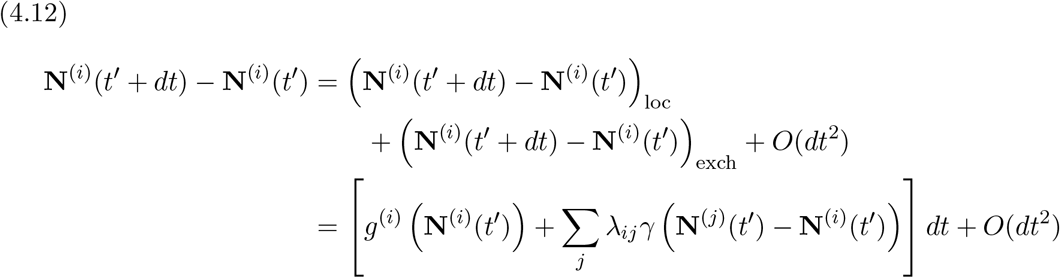

with arbitrarily high probability. Therefore, **N**^(*i*)^(*t*) is well-approximated by **Ñ**^(*i*)^(*t*).

As an example of the HFLSA, consider a two-host system in which each host has local dynamics (2.9). Let **N**^(1)^(0) = (2, 2) and **N**^(2)^(0) = (12, 12). In Figure 8, we show how the approximation improves as we increase the total-interaction-frequency parameter *λ*_tot_ = *λ*_12_ and decrease the interaction strength *γ* for fixed *λ*_tot_*γ* = 8. The error that we obtain by using the approximate microbiome abundance vector **Ñ** ^(*i*)^(*t*) appears to be largest near times that the *i*th abundance vector **Ñ**^(*i*)^(*t*) transitions between different basins of attraction. However, for sufficiently large *λ*_tot_, this error becomes arbitrarily small for all times *t* ∈ [0, *T*] with arbitrarily high probability.

**Figure 8.**
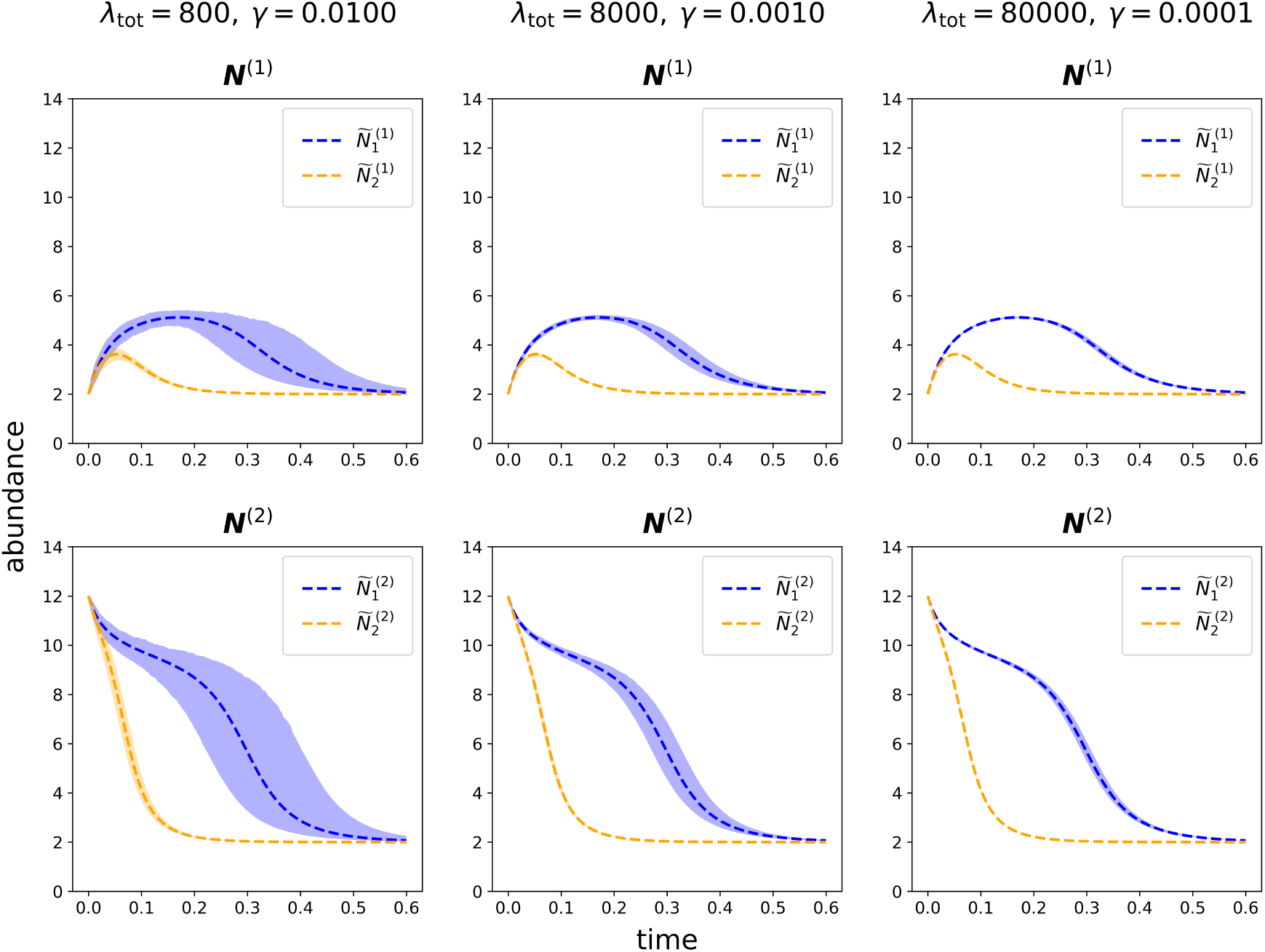
Numerical experiments for a two-host system in which each host has local dynamics (2.9). The three columns show experiments for different values of the total-interaction-frequency parameter λ_tot_ and the interaction strength γ for fixed λ_tot_γ = 8. We show the microbiome abundances for H^(1)^ in the first row and the microbiome abundances for H^(2)^ in the second row. We run 500 simulations for each set of parameters. The highlighted region shows the range between the 5^th^ and 95^th^ percentiles of the simulated host abundances. The dashed curves show the HFLSAs for these experiments.

### 4.2. High-Frequency, Constant-Strength Approximation (HFCSA)

The HFCSA is accurate when interactions are very frequent and have constant strengths. In this regime, all microbiome abundance vectors converge rapidly to the mean microbiome abundance vector

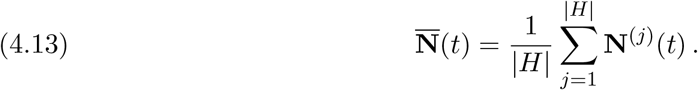

Subsequently, these “synchronized” microbiome abundance vectors each follow the mean of their local dynamics (see (4.17) below).

#### Theorem 4.2

(High-Frequency, Constant-Strength Approximation Theorem). *Fix the relative interaction-frequency parameters l*_*ij*_, *the interaction strength γ >* 0, *and a time T. Suppose that each local-dynamics function g*^(*i*)^ *is Lipschitz continuous and bounded (see* subsection 2.3*). Let ε* ∈ (0, 1], *δ >* 0, *and η >* 0 *be arbitrary but fixed constants. For sufficiently large λ*_tot_, *each host microbiome abundance vector* **N**^(*i*)^(*t*) *satisfies*

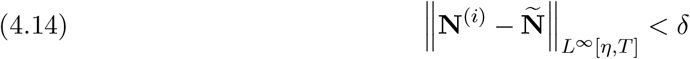

*with probability larger than* 1 − *ε, where*

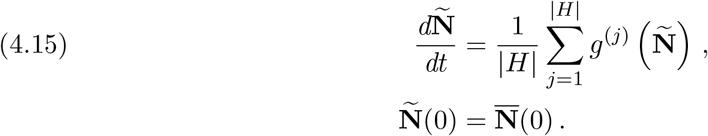

We provide key steps of the proof of Theorem 4.2 in this section. We give a full proof in Appendix C.

When two hosts *H*^(*i*)^ and *H*^(*j*)^ interact at time *t*_*I*_, they exchange portions of their microbiome as described in (2.1). This exchange causes no change in the sum

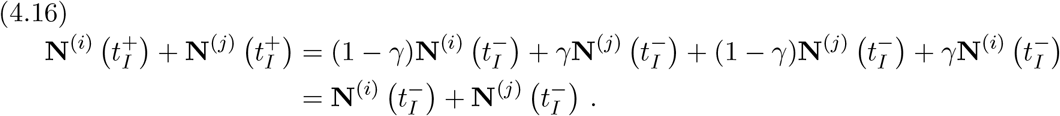

Therefore, interactions do not cause any direct change in the mean microbiome abundance vector 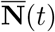. The local dynamics drive the evolution

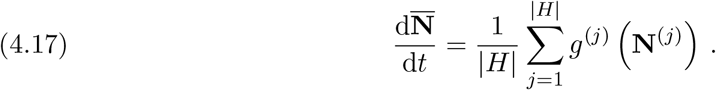

For sufficiently large *λ*_tot_, interactions occur on a much faster time scale than the local dynamics. Because of this separation of time scales, all microbiome abundance vectors **N**^(*i*)^(*t*) converge rapidly, which entails that

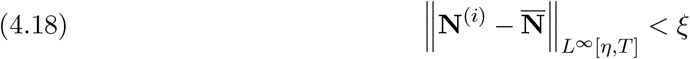

with high probability. For fixed probability, a larger *λ*_tot_ allows a smaller bound *ξ*. For a sufficiently small bound *ξ*, each abundance vector **N**^(*i*)^(*t*) is close enough to 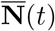 so that 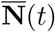 is well-approximated by the approximate microbiome abundance vector **Ñ** (*t*) (see (4.15)). For times *t* ∈ [*η, T*], each microbiome abundance vector **N**^(*i*)^(*t*) is very close to 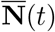 and 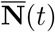 is very close to **Ñ** (*t*). Therefore, on the interval [*η, T*], each abundance vector **N**^(*i*)^(*t*) is well-approximated by **Ñ** (*t*).

As an example of the HFCSA, consider a two-host system in which each host has local dynamics (2.9). Let **N**^(1)^(0) = (2, 2) and **N**^(2)^(0) = (12, 12). In Figure 9, we show how the approximation improves as we increase the total-interaction-frequency parameter *λ*_tot_ = *λ*_12_ for fixed interaction strength *γ* = 0.02.

**Figure 9.**
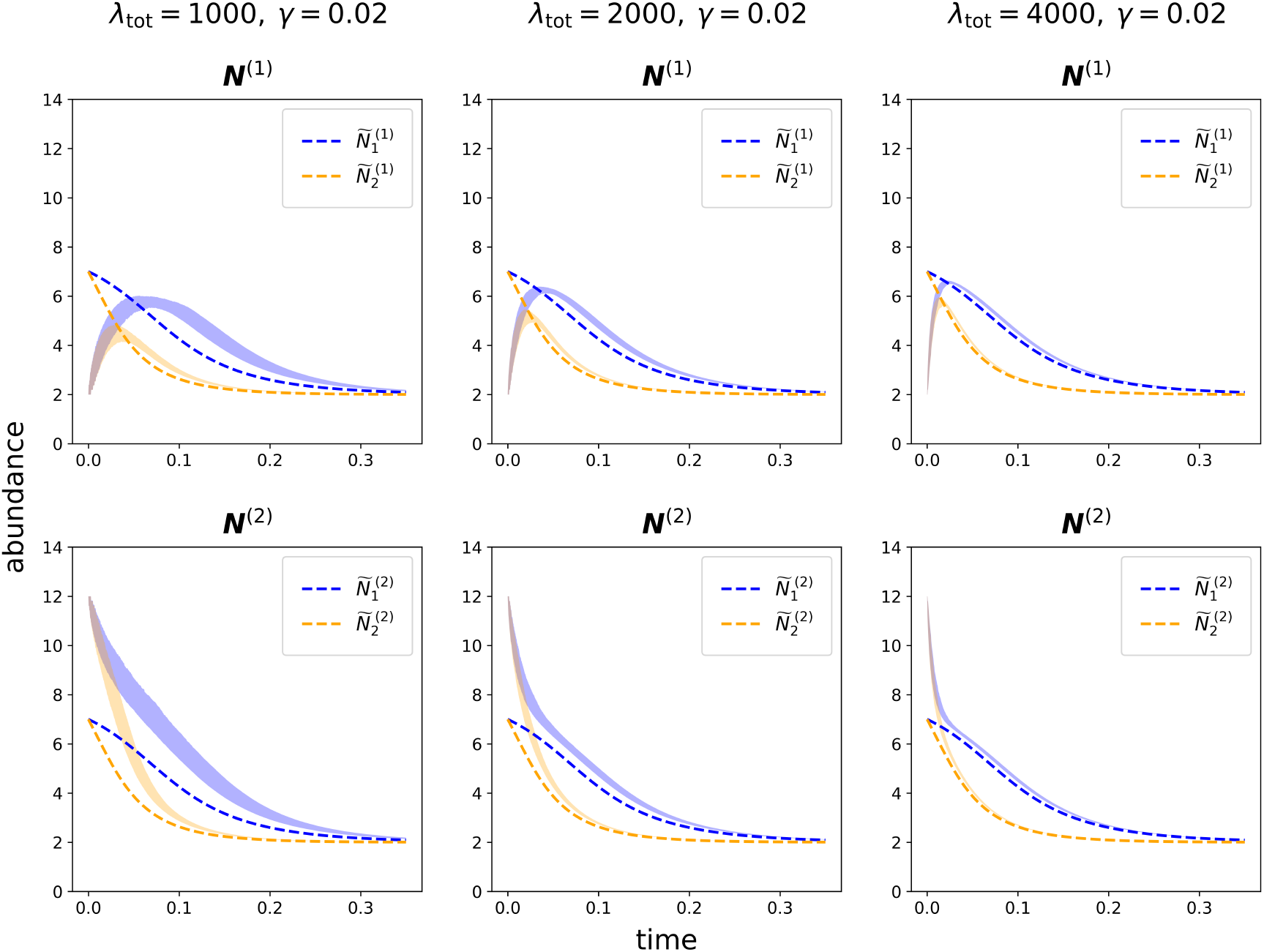
Numerical experiments for a two-host system in which each host has local dynamics (2.9). The three columns show experiments for different values of the total-interaction-frequency parameter λ_tot_ for fixed interaction strength γ = 0.02. We show the microbiome abundances for H^(1)^ in the first row and the microbiome abundances for H^(2)^ in the second row. We run 500 simulations for each set of parameters. The highlighted region shows the range between the 5^th^ and 95^th^ percentiles of the simulated host abundances. The dashed curves show the HFCSA for these experiments.

## 5. Numerical Experiments

In this section, we present simulations for a system of 10 hosts with local dynamics (2.9). We showed the interaction network for this system in Figure 1. This network has 25 edges, and we suppose that all relative interaction-frequency parameters *λ*_*ij*_ are equal. For each (*H*^(*i*)^, *H*^(*j*)^) ∈ *E*, the corresponding relative interaction-frequency parameter is *l*_*ij*_ = 1*/*25. We explore the accuracy of our three approximations for a range of values for the total-interaction-frequency parameter *λ*_tot_ and the interaction strength *γ*.

### 5.1. Pair Approximation for the LF

For the LFA, the approximate basin probability tensor 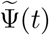 (see (3.9)) has dimension *m*_1_ *× m*_2_ *×* · · · *× m*_|*H*|_. In many circumstances, this tensor is too large to analyze directly. Therefore, we use a pair approximation [19, 20] of 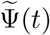 for our calculations of the LFA in this subsection.

We approximate 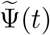 by tracking the individual probabilities

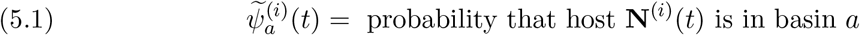

and the dyadic (i.e., pair) probabilities

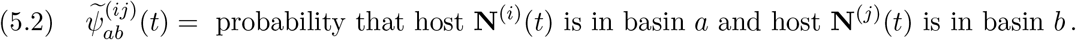

To obtain a pair approximation, we also need to consider the triadic (i.e., triplet) probabilities

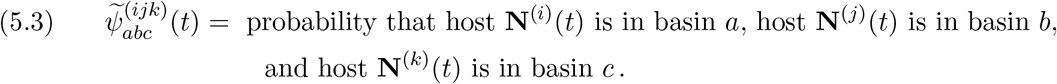

In the LFA, if an interaction between hosts *H*^(*i*)^ and *H*^(*j*)^ causes their microbiome abundance vectors to move from basin *d* to basin *a* and from basin *e* to basin *b*, respectively, then 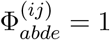. Otherwise, 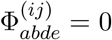. The probability that a given interaction is one between *H*^(*i*)^ and *H*^(*j*)^ is *l*_*ij*_. Therefore, the change in 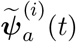 due to an interaction at time *t*_*I*_ is

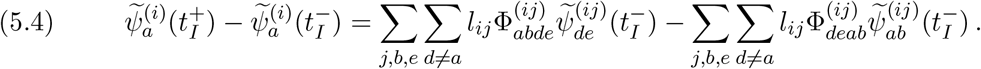

Using frequency-scaled time, the expected number of system interactions during an interval [*t*^*\**^, *t*^*\**^ + *dt*^*\**^] is *dt*^*\**^. Therefore,

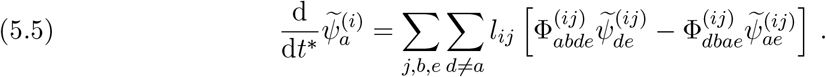

For the dyadic probabilities, we have

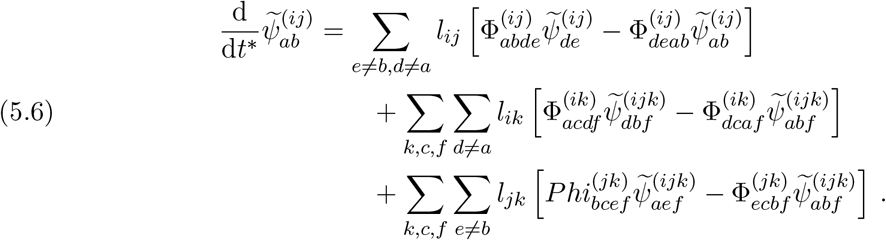

The right-hand sides in (5.5) and (5.6) are exact expressions. We form an approximation by replacing the triadic probabilities in (5.6) with combinations of dyadic probabilities. In the derivatives of 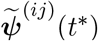, when considering the impact of interactions between hosts *H*^(*i*)^ and *H*^(*k*)^, we use the approximation

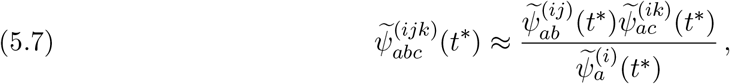

which assumes that 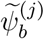 and 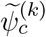 are independent. Inserting the approximation (5.7) into (5.6) gives

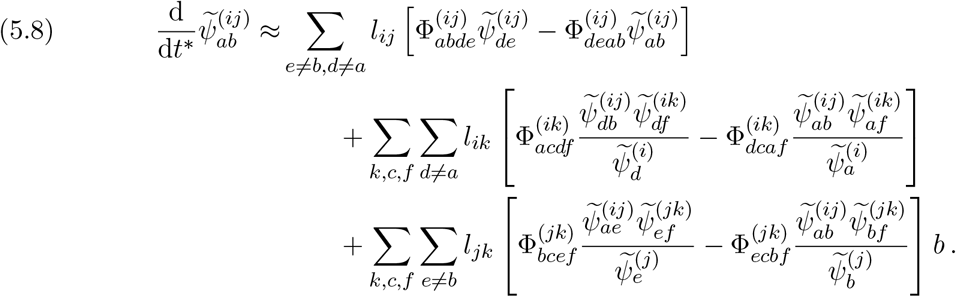

Equations (5.5) and (5.8) constitute a pair approximation of the evolution of 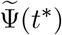. In our simulations (see subsection 5.2), we compare the individual probabilities 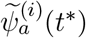 from our pair approximation (5.5, 5.8) with the fraction 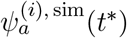 of simulations in which **N**^(*i*)^(*t*^*\**^) is in basin of attraction *a*.

### 5.2. Simulations for the Low-Frequency Approximation

In this subsection, we show numerical results for the LFA (3.9). We use the pair approximation (5.5, 5.8) to determine the

individual probabilities 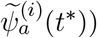. As we described in subsection 5.1, an individual probability 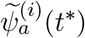 is an approximation of the probability from the LFA that **N**^(*i*)^(*t*^*\**^) is in basin of attraction *a*. We compare these individual probabilities to the fraction 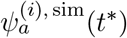 of simulations of (2.1, 2.9) in which **N**^(*i*)^(*t*^*\**^) is in basin of attraction *a*. We perform these approximations and simulations for 59 linearly spaced interaction strengths *γ* ∈ [0, 0.5] and 13 logarithmically spaced values of the total-interaction-frequency parameter *λ*_tot_ ∈ [2.5 *×* 10^−2^, 2.5 *×* 10^4^]. Because ℬ = {0.1, 0.4} for this system, the LFA is not valid when *γ* = 0.1 or *γ* = 0.4, so we exclude these values of *γ* from our simulations. Therefore, we perform simulations for values of *γ* ∈ [0, 0.5] that are multiples of 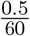 (except *γ* = 0.1 and *γ* = 0.4). For each pair of *γ* and *λ*_tot_ values, we select a random four-dimensional (4D) vector 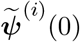 from the Dirichlet distribution Dir(1, 1, 1, 1) [14] for each host. The entries of each of these 4D vectors sum to 1. For each simulation, we set each **N**^(*i*)^(0) to be the stable equilibrium point in one of the basins of attraction. For each basin of attraction *a*, the initial microbiome abundance vector **N**^(*i*)^(0) is in that basin of attraction with probability 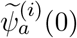.

We perform 1000 simulations of (2.1, 2.9) for each pair of *γ* and *λ*_tot_ values, and we calculate the fraction 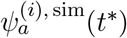 of the simulations in which **N**^(*i*)^(*t*^*\**^) is in basin of attraction *a*. We compare 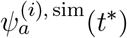 to 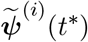, which we calculate using the pair approximation (5.5, 5.8). We calculate 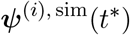 and 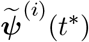 for 1001 evenly spaced frequency-scaled times 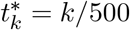 in the interval [0, 2]. In Figure 10, we plot the error

**Figure 10.**
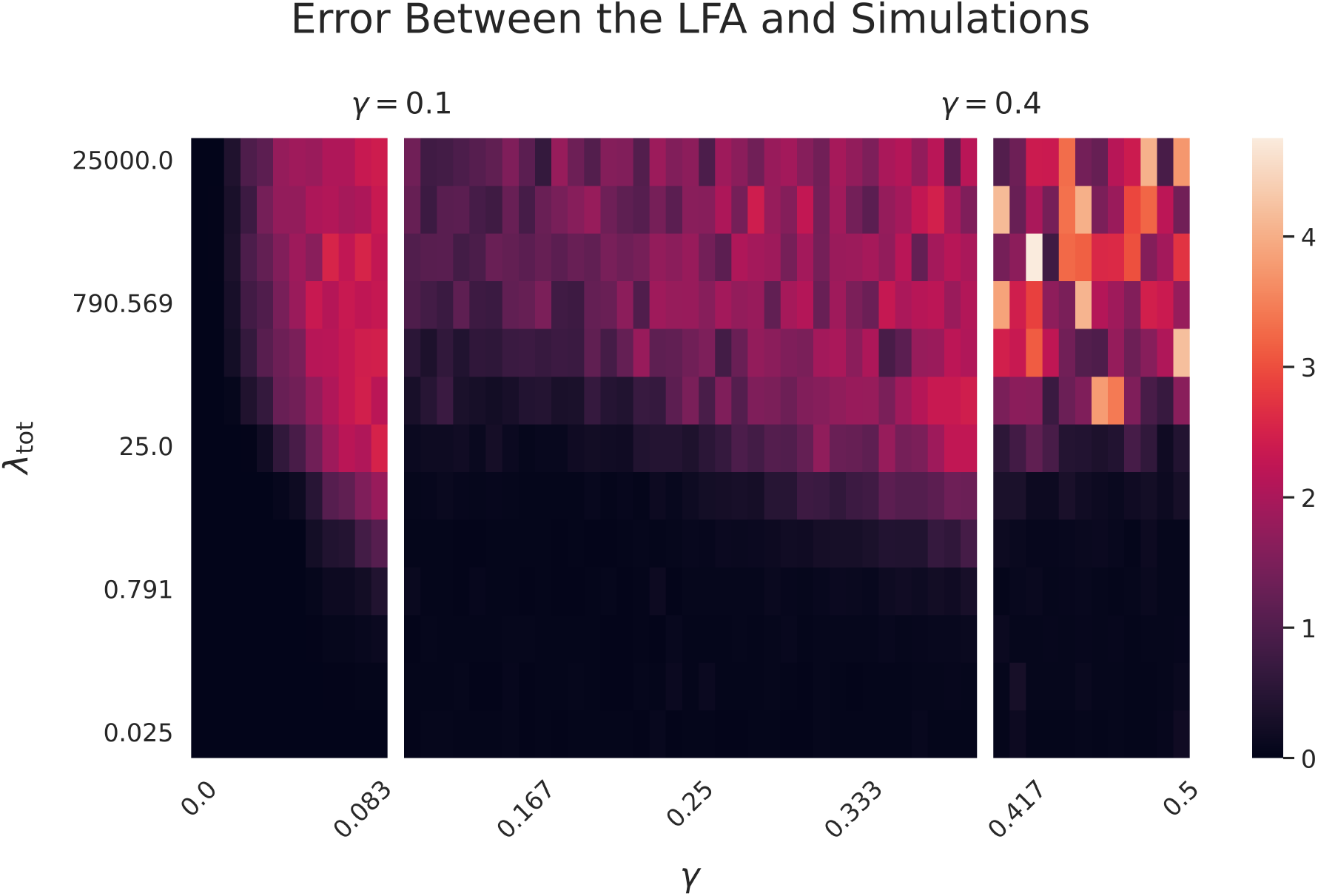
The error (5.9) between the LFA (see (3.9)) versus means of 1000 simulations of (2.1, 2.9) for each pair of the interaction strength γ and the total-interaction-frequency parameter λ_tot_. We plot γ on a linear scale and λ_tot_ on a logarithmic scale. We do not plot errors for γ = 0.1 and γ = 0.4 because the LFA is not valid for these values.

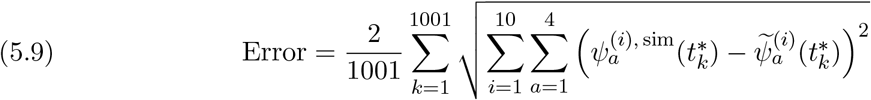

for each pair of *γ* and *λ*_tot_ values. The error (5.9) is a discrete approximation of the norm 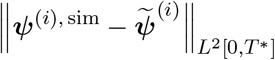.

The LFA is most accurate when the total-interaction-frequency parameter *λ*_tot_ is small. It is also better when the interaction strength *γ* is not near 0.1 or 0.4. There are two types of errors in the LFA. The first type of error arises when repeated interactions occur in sufficiently quick succession to yield a transition that the LFA misses. For example, for *γ ≈*0.342, the LFA does not predict that the **N**^(*i*)^(*t*) can move from basin 2 to basin 1. For sufficiently large values of *λ*_tot_, repeated interactions in short succession are common (and not merely possible), which causes the LFA to overestimate the probability that a host is in basin 2 and underestimate the probability that a host is in basin 1. In Figure 11, we illustrate this type of error for *γ ≈* 0.342 and several values of *λ*_tot_.

**Figure 11.**
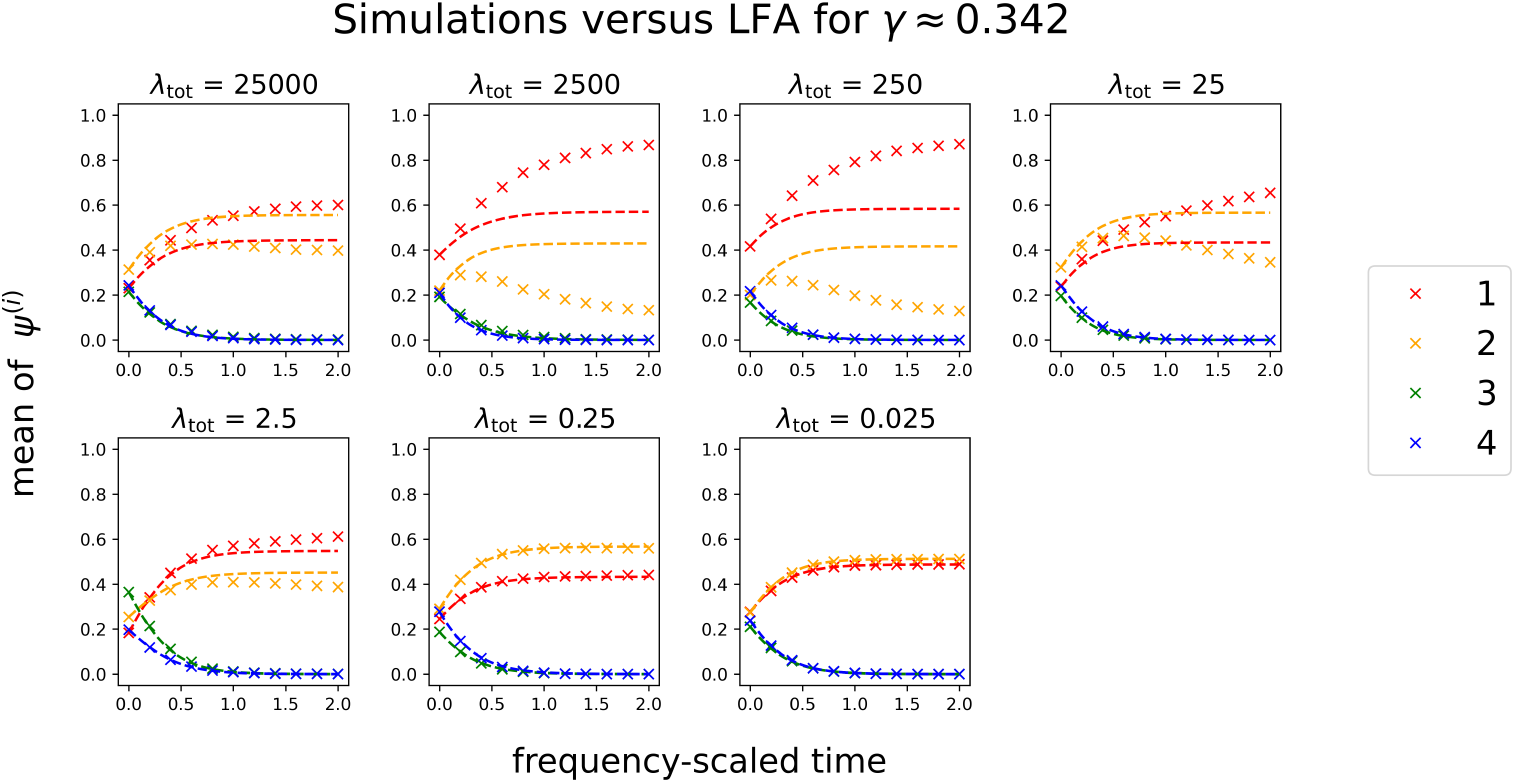
The means of the simulated probabilities 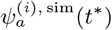 over all hosts for interaction strength γ ≈ 0.342 and several values of the total-interaction-frequency parameter λ_tot_. The dashed curves indicate the LFA approximation of the mean of the probabilities 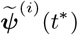 over all hosts.

The second type of error arises when repeated interactions in sufficiently quick succession cause the LFA to overestimate the impact of the second and subsequent interactions. As an example, for the interaction strength *γ ≈* 0.433, the LFA predicts that **N**^(*j*)^(*t*) will be in basin 1 after an interaction whenever *H*^(*j*)^ interacts with *H*^(*i*)^ and **N**^(*i*)^(*t*) is in basin 1 before the interaction. This prediction arises because the LFA assumes that **N**^(*i*)^ = (2, 2) before this interaction. However, if *H*^(*i*)^ recently interacted with a different host, then **N**^(*i*)^(*t*) may be in basin 1 but not sufficiently close to the equilibrium point (2, 2) to drive a transition to the basin of attraction of **N**^(*j*)^(*t*). Consequently, the LFA overestimates the probability that a host is in basin 1. In Figure 12, we illustrate this type of error for *γ ≈* 0.433 and several values of *λ*_tot_.

**Figure 12.**
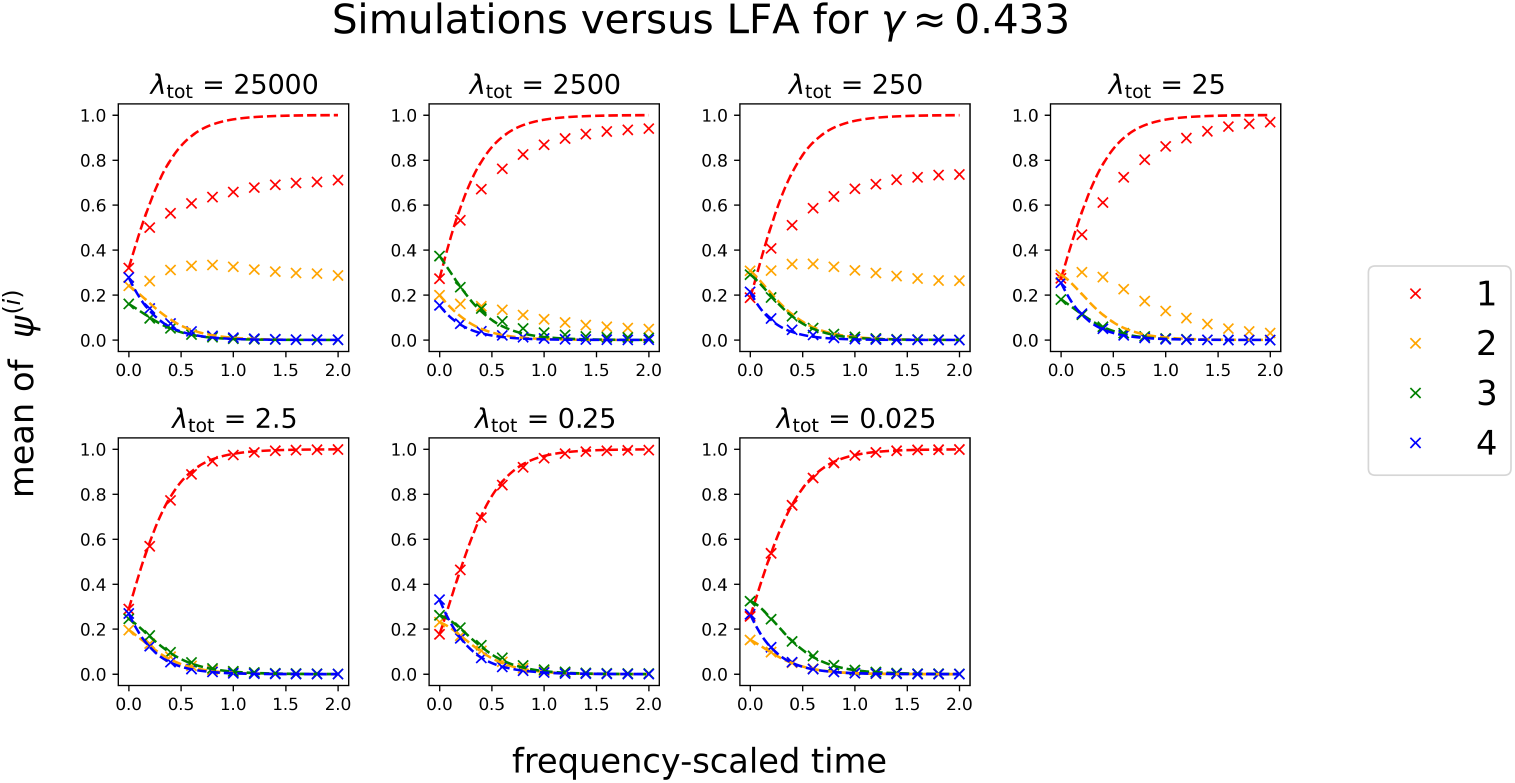
The mean of the simulated probabilities 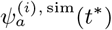 over all hosts for interaction strength γ ≈ 0.433 and several values of the total-interaction-frequency parameter λ_tot_. The dashed curves indicate the LFA approximation of the mean of the probabilities 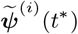 over all hosts.

### 5.3. Simulations for the High-Frequency Approximations

In this subsection, we show numerical results for the HFLSA (4.2) and the HFCSA (4.15). We compare the approximate microbiome abundance vectors from these two approximations to simulations of (2.1, 2.9).

To evaluate the accuracy of the approximate microbiome abundance vectors **Ñ**^(*i*)^(*t*) that we obtain from the HFLSA (4.2), we compare them to the microbiome abundance vectors **N**^(*i*)^(*t*) from (2.1, 2.9). We perform these simulations for 13 logarithmically spaced values of the total-interaction-frequency parameter *λ*_tot_ ∈ [25, 2500] and 75 linearly spaced values of *λ*_tot_*γ* ∈ [0.04, 0.3]. We use the product *λ*_tot_*γ* as a parameter instead of the interaction strength *γ* on its own to illustrate the improvement of the HFLSA as we increase *λ*_tot_ for fixed *λ*_tot_*γ*. For each pair of *λ*_tot_ and *λ*_tot_*γ* values, we perform 1000 simulations over the time interval [0, 1] with initial conditions

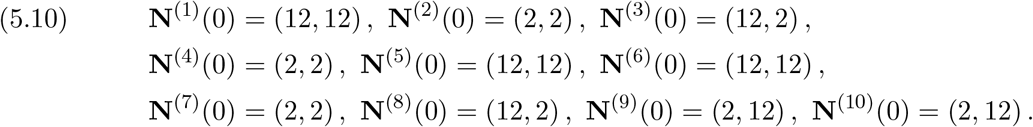

The microbiome abundance vector **N**^(*i*),*l*^(*t*) is the *l*th simulated microbiome abundance vector for host *H*^(*i*)^. For each of these simulations, we calculate the simulated microbiome abundance vectors **N**^(*i*),*l*^(*t*) for 101 evenly spaced times *t*_*k*_ = *k/*100. We compare these simulations to the approximate microbiome abundance vector **Ñ**^(*i*)^(*t*) from the HFLSA (4.2). In Figure 13, we plot the error

**Figure 13.**
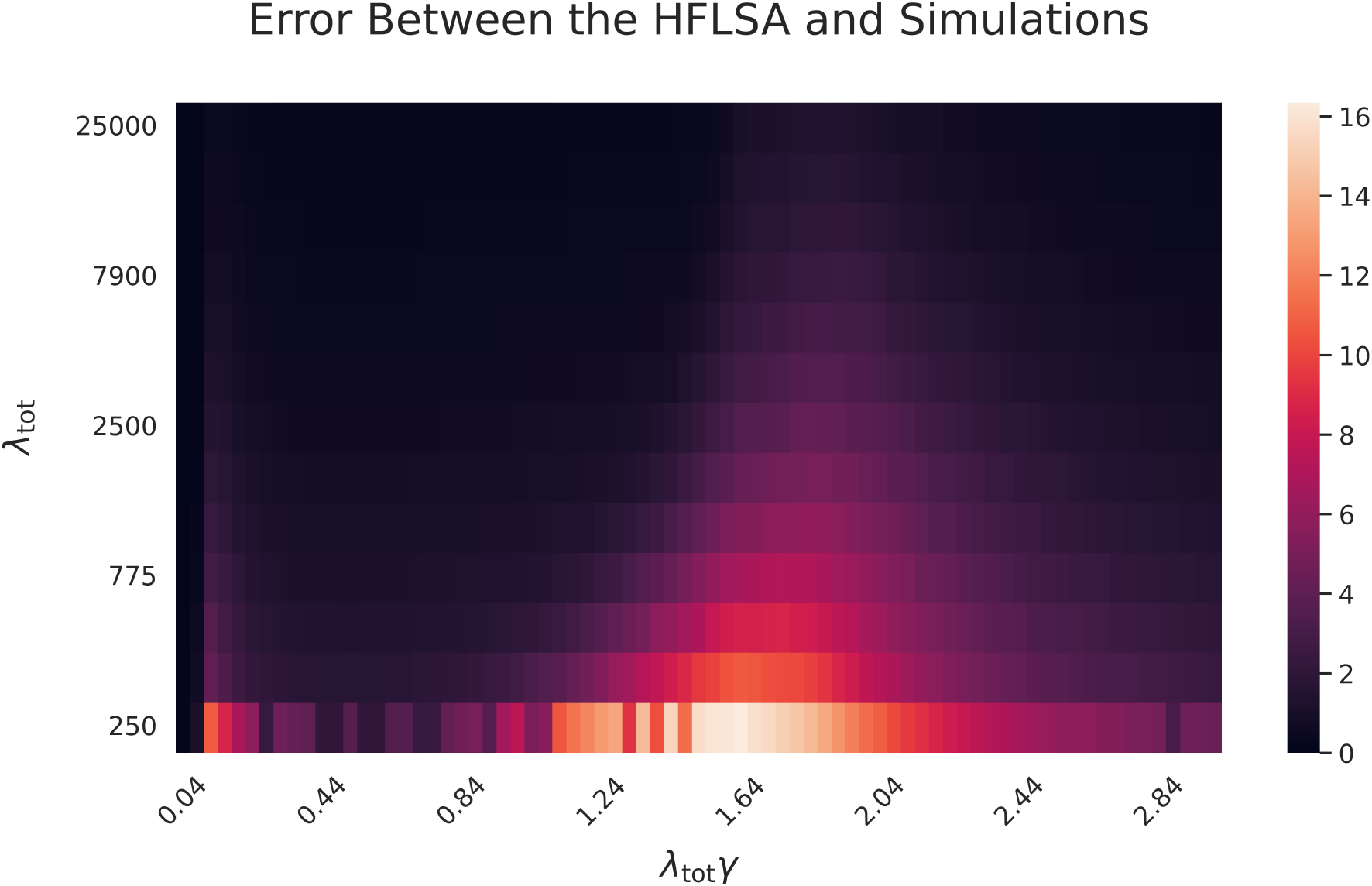
The mean error (5.11) between the approximate microbiome abundance vectors {**Ñ**^(i)^(t)} from the HFLSA (see (4.2)) and the microbiome abundance vectors **N**^(i)^(t) for 1000 simulations of (2.1, 2.9). We plot λ_tot_ on a logarithmic scale and plot λ_tot_γ on a linear scale.

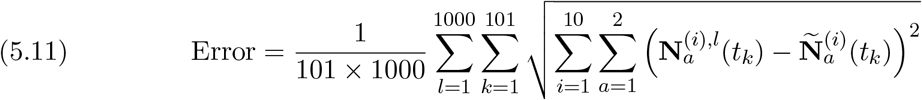

for each pair of *λ*_tot_ and *λ*_tot_*γ* values. The error (5.11) is a discrete approximation of the mean of 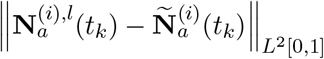 over all simulations. For any fixed value of *λ*_tot_ *γ*, the HFLSA is more accurate for larger *λ*_tot_. However, for a fixed value of *λ*_tot_, the error depends significantly on the value of *λ*_tot_*γ*. Each value of *λ*_tot_*γ* yields a set {**Ñ**^(*i*)^(1)} of final approximate microbiome abundance vectors. For all but a finite set of values of *λ*_tot_*γ*, each final approximate microbiome abundance vector **Ñ**^(*i*)^(1) changes continuously with *λ*_tot_*γ*. However, there are a finite number of *λ*_tot_*γ* values for which some final approximate microbiome abundance vector **Ñ**^(*i*)^(1) has a discontinuous jump. For the initial set of microbiome abundance vectors (5.10), these discontinuous jumps occur at

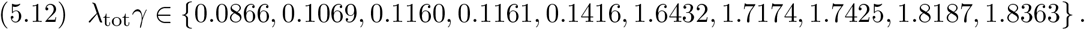

The regions in which the HFLSA performs worst in our simulations are near *λ*_tot_*γ ≈* 0.1 and *λ*_tot_*γ ≈* 1.8, which are very close to several of the values in (5.12).

For the HFCSA (4.15), we perform simulations for 61 linearly spaced values of the interaction strength *γ* ∈ [0, 0.5] and 13 logarithmically spaced values of the total-interaction-frequency parameter *λ*_tot_ ∈ [2.5 *×* 10^−1^, 2.5 *×* 10^4^]. We use the same initial conditions (5.10) as in our HFLSA simulations. We also again perform 1000 simulations and evaluate each simulated microbiome abundance vector **N**^(*i*),*l*^(*t*) for 101 evenly spaced times *t*_*k*_ = *k/*100 on the time interval [0, 1]. In Figure 14, we show the error (5.11) for each pair of *γ* and *λ*_tot_ values. The approximation is accurate for sufficiently large *λ*_tot_. In general, a larger *γ* increases the rate at which each microbiome abundance vector **N**^(*i*)^(*t*) converges to the mean microbiome abundance vector 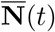. Consequently, the HFCSA yields a better approximation of **N**^(*i*)^(*t*) for larger values of *γ*.

**Figure 14.**
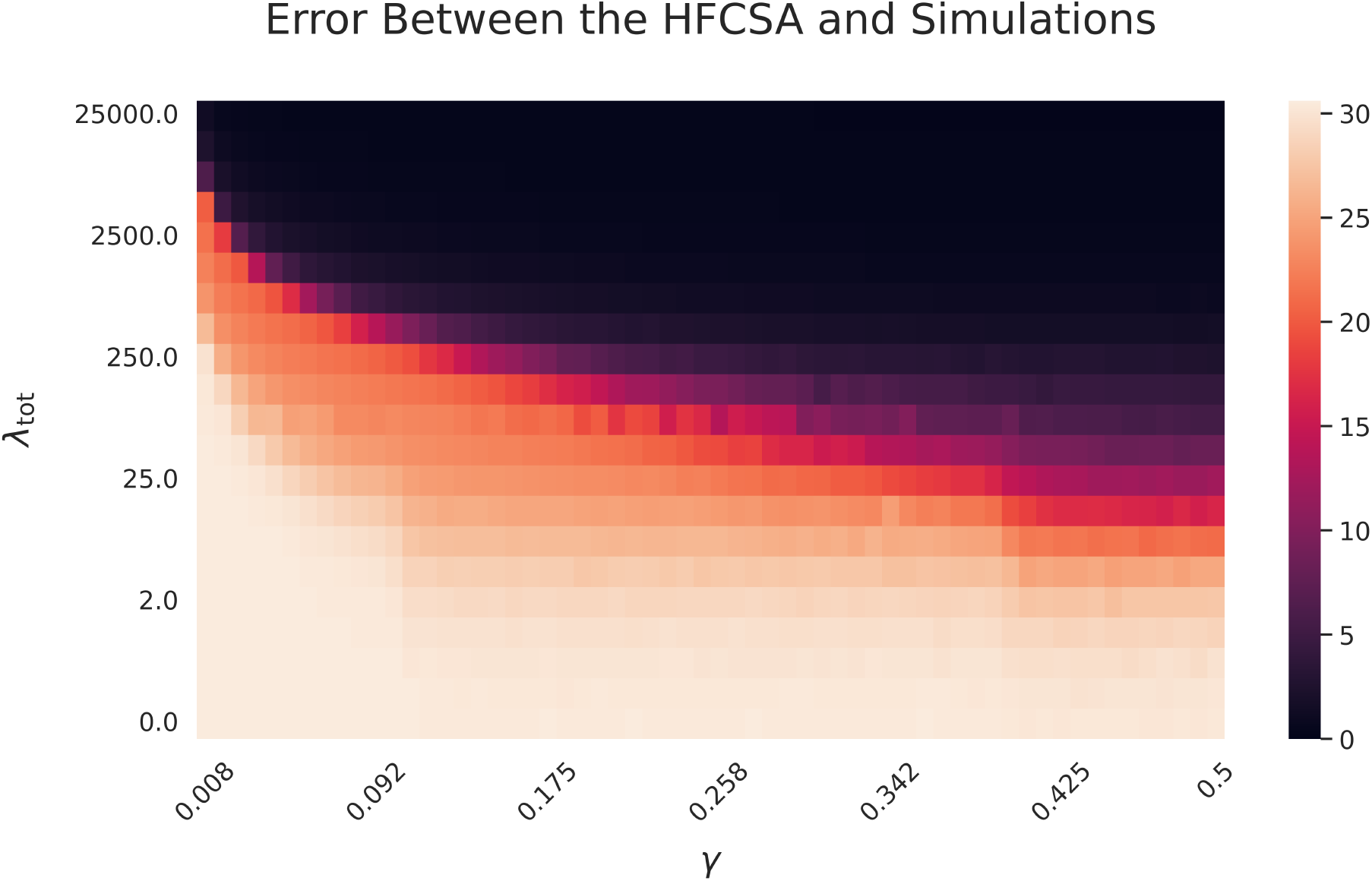
The mean error (5.11) between the approximate microbiome abundance vectors {**Ñ**^(i)^(t)} from the HFCSA (see (4.15)) and the microbiome abundance vectors **N**^(i)^(t) for 1000 simulations of (2.1, 2.9). We plot λ_tot_ on a logarithmic scale and plot γ on a linear scale.

## 6. Conclusions and Discussion

### 6.1. Summary

We developed a novel framework to model the microbiome dynamics of living hosts that incorporates both the local dynamics within an environment and exchanges of microbiomes between environments. Our framework extends existing metacommunity theory by accounting for the discrete nature of host interactions. Unlike classical mass-effects models, our framework incorporates two distinct parameters that control interaction frequencies and interaction strength. Using both analytical approximations and numerical computations, we demonstrated that both parameters are necessary to determine microbiome dynamics.

We developed approximations in three parameter regions, and we proved their accuracy in those regions. Our low-frequency approximation (LFA) gives a good approximation of the microbiome dynamics when local dynamics are much faster than host interactions. Our high-frequency, low-strength approximation (HFLSA) encodes the dynamics of a system when interactions are frequent but weak, resulting in a model with the same form as the mass-effects model (1.3). Finally, our high-frequency, constant-strength approximation (HFCSA) accurately predicts the rapid convergence of all hosts’ microbiome dynamics when interactions are frequent and have constant interaction strength *γ*. We validated each of these approximations through numerical experiments on an illustrative model of microbiome dynamics for a range of parameter values.

A qualitative example of dynamics in our model involves the probability that all microbiome abundance vectors converge in some time interval. This probability depends both the interaction-frequency parameters and on the interaction strength. Using the LFA, we showed for sufficiently small interaction-frequency parameters that the convergence probability depends on whether the interaction strength *γ* is large enough that a single interaction between two hosts places their microbiome abundance vectors in the same basin of attraction. By contrast, using the HFLSA, we showed for sufficiently large interaction-frequency parameters that the convergence probability depends on the product *λ*_tot_*γ* of the total-interaction-frequency parameter *λ*_tot_ and the interaction strength *γ*. For intermediate values of the interaction-frequency parameters, we used numerical simulations of models in our framework to examine convergence probabilities.

### 6.2. Outlook

Our modeling framework provides a foundation for many promising future research directions in microbiome dynamics. In our framework’s current form, one can use it to study the effects of host interactions in many ecological models of local dynamics. One can also use our framework to study the impact of the structure of interaction networks on microbiome dynamics.

There are many possible extensions of our modeling framework. For example, we considered a homogeneous interaction strength *γ* for simplicity. However, one can allow each pair of hosts to have heterogeneous interaction strengths *γ*_*ij*_. Additionally, hosts can exchange different microbe species with different exchange strengths, so one can also consider interaction strengths *γ*_*ijk*_ that encode the exchange of different species *k* when hosts *H*^(*i*)^ and *H*^(*j*)^ interact. This seems helpful when using consumer–resource models for the local dynamics. In this case, it also may be desirable to separately encode the strengths of microbiome exchange and resource exchange. Another way to extend our framework is to relax our assumption of instantaneous microbiome exchange by instead employing rapid but continuous functions.

Additionally, in our framework, the LFA assumes that the attractors of each local-dynamics function *g*^(*i*)^ consist of a finite set of stable equilibrium points. We believe that it is possible to extend the LFA to systems in which the *g*^(*i*)^ have more complicated attractors, such as limit cycles and chaotic attractors. We also believe that it is possible to generalize the HFLSA and HFCSA to systems in which the times between consecutive interactions for pairs of adjacent hosts follow a distribution other than an exponential distribution.

## Appendix A. Proof of Low-Frequency Approximation Theorem

In this appendix, we prove the LFA Theorem (see Theorem 3.1).

### Theorem A.1

(Low-Frequency-Approximation Theorem). *Suppose that the attractors of each host’s local dynamics consist of a finite set of stable equilibrium points at which the local-dynamics function g*^(*i*)^ *is inward pointing, and let each g*^(*i*)^ *be continuous and bounded (see* subsection 2.3*). Fix γ*∉ ℬ, *all l*_*ij*_, *and a frequency-scaled time T*^*\**^. *As λ*_tot_ *→* 0, *the basin probability tensor* Ψ(*t*^*\**^) *converges uniformly to* 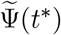 *on* [0, *T*^*\**^], *where*

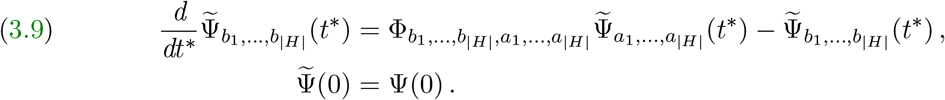

*Proof*. Each *g*^(*i*)^ is bounded (see subsection 2.3) so, there exists a constant *M* such that each entry of **N**^(*i*)^(*t*) is nonnegative and

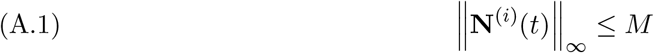

for each microbiome abundance vector **N**^(*i*)^(*t*) and all times *t ≥* 0.

Consider an arbitrary but fixed *ε >* 0. We will show that

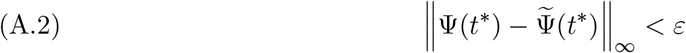

for sufficiently small total-interaction-frequency parameter *λ*_tot_ and all frequency-scaled times *t*^*\**^ ∈ [0, *T*^*\**^].

For each *i*, let 𝒜_*i*_ be the set of stable equilibrium points of the local dynamics of host *H*^(*i*)^. Suppose that adjacent hosts *H*^(*i*)^ and *H*^(*j*)^ interact at time *t*_*I*_ and that 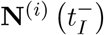 and 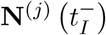 are at stable equilibrium points **a**^(*i*)^ ∈ 𝒜_*i*_ and **a**^(*j*)^ ∈ 𝒜_*j*_, respectively. For **x, y** ∈ [0, *M*]^*n*^, let

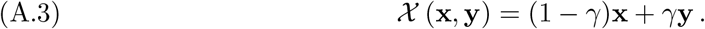

After the interaction, 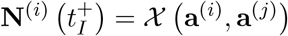 and 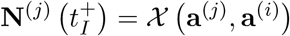. Because *γ*∉ ℬ,

it follows that 𝒳 **a**^(*i*)^, **a**^(*j*)^ and 𝒳 **a**^(*j*)^, **a**^(*i*)^ are in some basins of attraction *b*_*i*_ and *b*_*j*_ for the stable equilibrium points **b**^(*i*)^ ∈ 𝒜_*i*_ and **b**^(*j*)^ ∈ 𝒜_*j*_, respectively.

Let

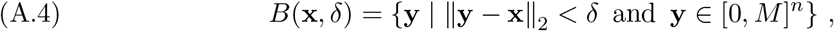

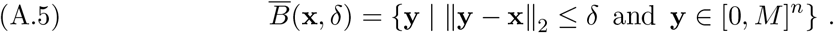

The basins of attraction of stable equilibrium points are open sets, so there exists *δ*(**a**^(*i*)^, **a**^(*j*)^) = *δ*(**a**^(*j*)^, **a**^(*i*)^) such that

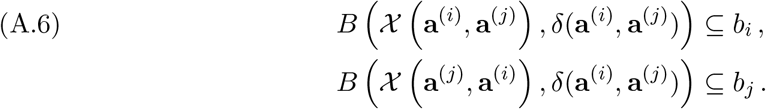

All 𝒜_*i*_ are finite, so there are minima

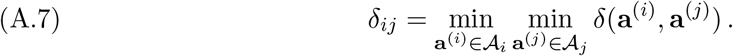

Let the flow **X**^(*i*)^(*t*, **x**) be the solution of

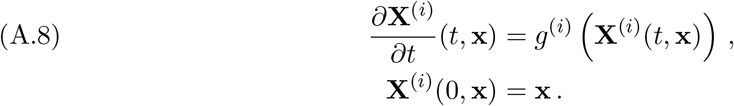

For each local-dynamics function *g*^(*i*)^, each **a**^(*i*)^ ∈ 𝒜_*i*_ is a stable equilibrium point at which *g* is inward pointing. Therefore, there exists *δ* **a** such that *g* (**y**) **a y** *>* 0 for 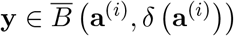. If **x** is in the basin of attraction of **a**^(*i*)^, then **X**(*t*_*a*_, **x**) ∈ *B* **a**^(*i*)^, *δ* **a**^(*i*)^ for some time *t*_*a*_. We then have

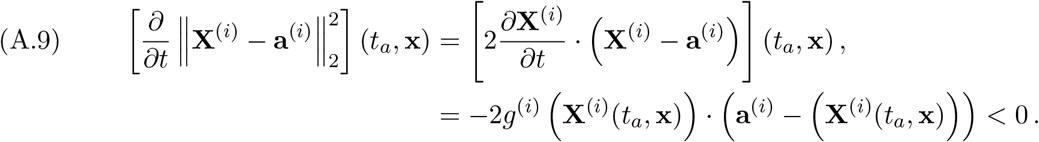

Consequently, ‖**X**^(*i*)^(*t*, **x**) − **a**^(*i*)^ ‖_2_ is monotonically decreasing for *t ≥ t*_*a*_. There are only finitely many hosts, so there exists

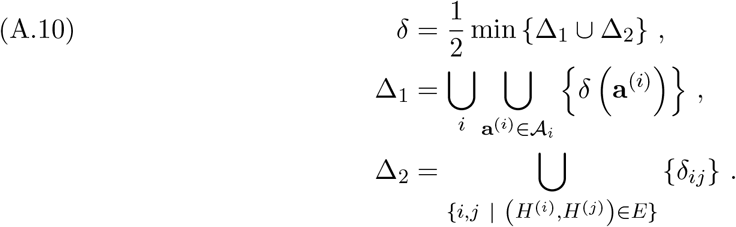

For any **x, y** ∈ [0, *M*]^*n*^, let

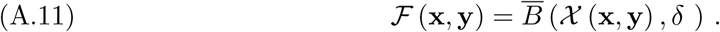

For each pair of adjacent hosts *H*^(*i*)^ and *H*^(*j*)^ and each **a**^(*i*)^ ∈ 𝒜_*i*_ and **a**^(*j*)^ ∈ 𝒜_*j*_, we have

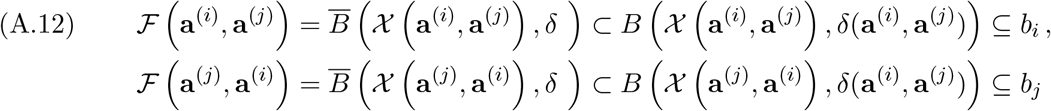

for some basins of attraction *b*_*i*_ and *b*_*j*_ of the local dynamics of hosts *H*^(*i*)^ and *H*^(*j*)^, respectively.

Recall that 𝒰^(*i*)^ ∈ [0, *M*]^*n*^ is the set of points that are not in the basin of attraction of some stable equilibrium point of the local dynamics of host *H*^(*i*)^, and let

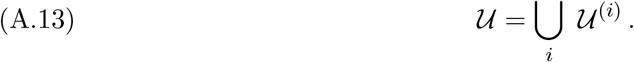

Each 𝒰^(*i*)^ has measure 0, so 𝒰 also has measure 0. Let **x** ∈ [0, *M*]^*n*^ *\* 𝒰. For each *i*, the vector **x** is i n the basin of att raction of some **a**^(*i*)^ ∈ *A*_*i*_. Therefore, there exists some time *t*_*a*_ such that ‖**X**^(*i*)^(*t*_*a*_, **x**) − **a**^(*i*)^ ‖_2_ = *δ*. We refer to such a time as a *crossing time*. Moreover, because 2*δ ≤ δ* (**a**^(*i*)^), it follows from (A.9) that ‖ **X**^(*i*)^(*t*, **x**) − **a**^(*i*)^ ‖_2_ is monotonically decreasing in *t* on some interval (*t*_*a*_ − *η, t*_*a*_ + *η*) for all *t ≥ t*_*a*_. T herefore, the crossing time *t*_*a*_ is the unique time that satisfies the equality ‖**X**^(*i*)^(*t*_*a*_, **x**) − **a**^(*i*)^ = *δ*‖_2_. Because the crossing time is unique, we

can define a function 𝒯 ^(*i*)^ (**x**) such that ‖ **X**^(*i*)^ (𝒯^(*i*)^ (**x**), **x**)− **a**(*i*) ‖ _2_ = *δ*. This function 𝒯^(*i*)^ (**x**) gives the unique crossing time for a flow that starts at **x**.

The local-dynamics function *g*^(*i*)^ is continuous, so

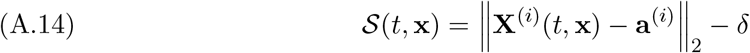

is continuously differentiable with respect to both *t* and **x**. Evaluating *S*(*t*, **x**) at *t* = 𝒯 ^(*i*)^(**x**) yields 𝒮 (𝒯 ^(*i*)^(**x**), **x**) = 0. The function 𝒮 (*t*, **x**) is monotonically decreasing in *t* on some interval (𝒯 ^(*i*)^(**x**) − *η*, 𝒯 ^(*i*)^(**x**) + *η*). Therefore, by the Implicit Function Theorem, 𝒯 ^(*i*)^ is a continuous function on each basin of attraction of *g*^(*i*)^.

Because 𝒯 ^(*i*)^(**x**) is continuous and each set *F*(**a**^(*i*)^, **a**^(*j*)^) is compact, there exist times

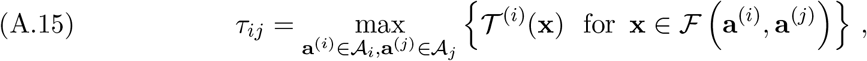

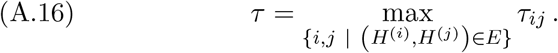

Suppose that an interaction occurs between hosts *H*^(*i*)^ and *H*^(*j*)^ at time *t*_*I*,1_. Let

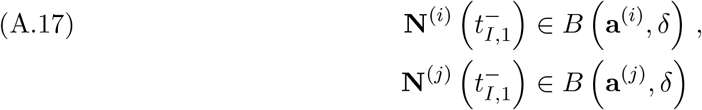

for some **a**^(*i*)^ ∈ 𝒜_*i*_ and **a**^(*j*)^ ∈ 𝒜_*j*_. We then have

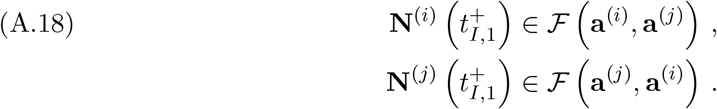

If the next interaction occurs at time *t*_*I*,2_ *> t*_*I*,1_ + *τ*, then

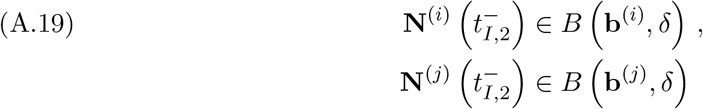

for some **b**^(*i*)^ ∈ 𝒜_*i*_ and **b**^(*j*)^ ∈ 𝒜_*j*_. If no two interactions occur within time *τ* of each other, then

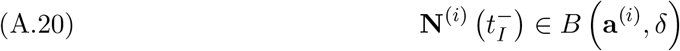

for all *i*, all interaction times *t*_*I*_, and some **a**^(*i*)^ ∈ 𝒜_*i*_. In this case, the effect of each interaction on the basin probability tensor Ψ is described exactly by the operation of the interaction operator

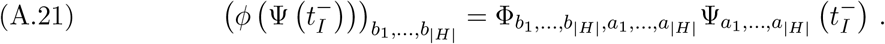

Let ℐ be the set of all possible sets 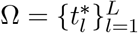 of frequency-scaled interaction times in the interval [0, *T*^*\**^]. We select a set Ω of interaction times using a Poisson process on [0, *T*^*\**^] with rate parameter 1. Let *q* : *I → R*^+^ be the probability density function for these interactions. Therefore,

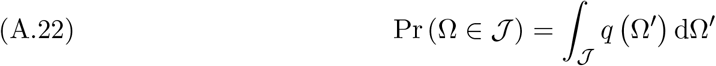

for any 𝒥 *⊆* ℐ. We define a counting function

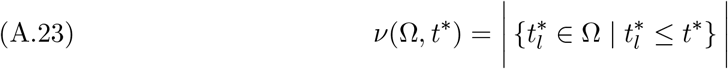

that tracks the number of interactions that occur in the interval [0, *t*^*\**^] for the set Ω. The approximate basin probability tensor 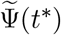 from the LFA (see (3.9)) conditioned on Ω is

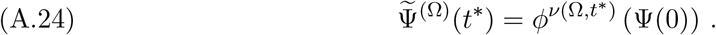

Therefore, (A.25)

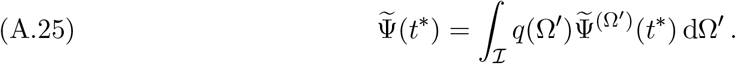

Let Ψ^(Ω)^(*t*^*\**^) be the basin probability tensor conditioned on Ω. We then have

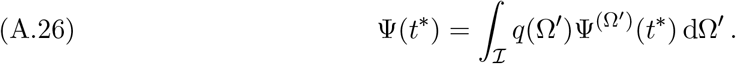

If 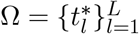 satisfies

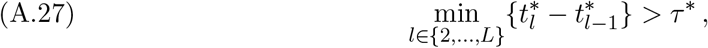

Then

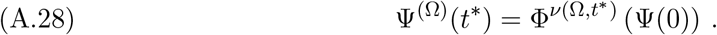

Let ℐ_1_ *⊂* ℐ be the set of interaction sets 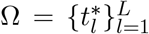 for which (A.27) holds, and let ℐ_2_ = ℐ \ ℐ_1_ *⊂* ℐ. Choose *dt*^*\**^ such that *τ*^*\**^ *< dt*^*\**^ *<* 2*τ*^*\**^ and *T*^*\**^*/dt*^*\**^ is an integer. For sufficiently small *τ*^*\**^, this is always possible. If two interaction times 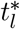 and 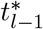 occur within *τ*^*\**^ of each other, then both interactions occur in an interval [*k dt*^*\**^, (*k* − 1) *dt*^*\**^] and/or an interval 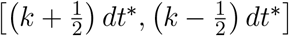 for an integer *k*. (These two types of intervals overlap.) All of these intervals have width *dt*^*\**^. Therefore, the probability that at least two interactions occur in a specific one of these intervals is

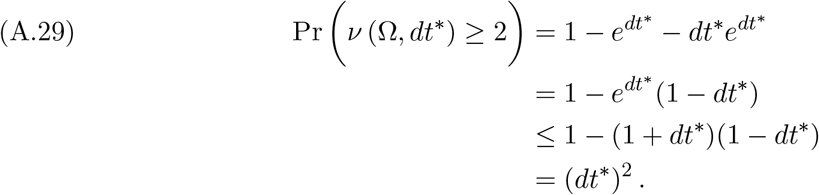

There are 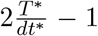 such intervals. Therefore, the probability that at least two interactions occur in at least one of these intervals is less than 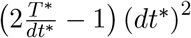. This probability equals the probability that a set Ω of interactions satisfies Ω ∈ ℐ_2_. Consequently,

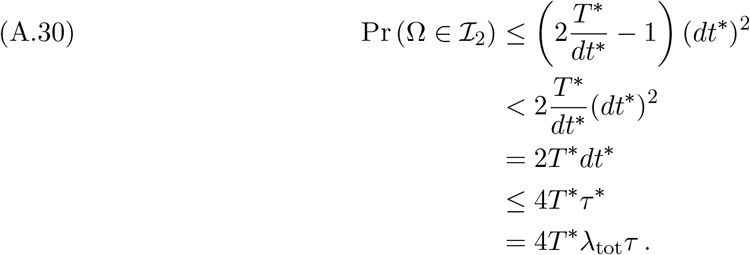

We choose 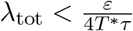, and we then have

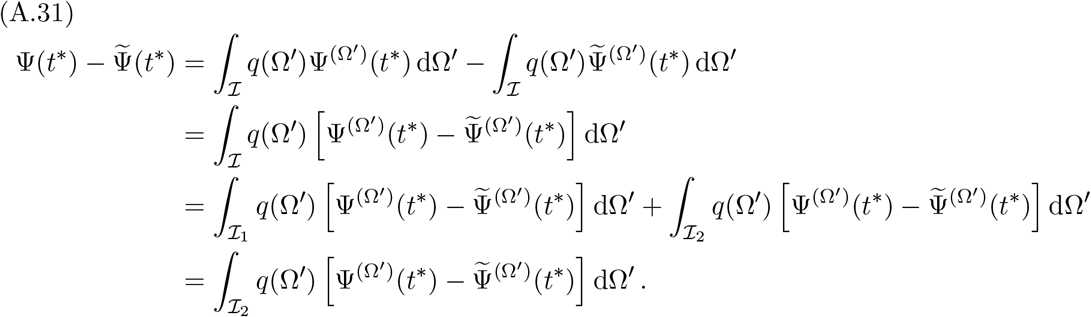

Therefore,

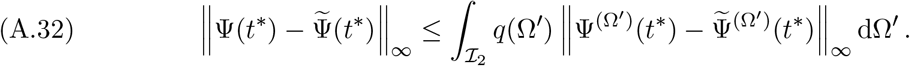

Each entry of Ψ(*t*^*\**^) and 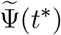 is a probability and hence is in the interval [0, 1], so we know that 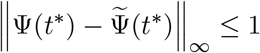. Therefore,

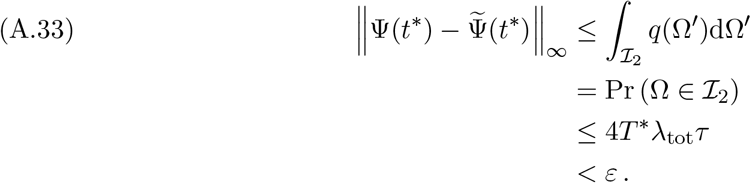

This bound holds for all *t*^*\**^ ∈ [0, *T*^*\**^]. Because *ε >* 0 is arbitrary, the basin probability tensor Ψ(*t*) converges uniformly to 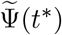 on [0, *T*^*\**^] as *λ*_tot_ *→* 0.

## Appendix B. Proof of High-Frequency Low-Strength Approximation Theorem

In this appendix, we prove the HFLSA Theorem (see Theorem 4.1).

### Theorem B.1

(High-Frequency, Low-Strength Approximation Theorem). *Fix the relative interaction-frequency parameters l*_*ij*_, *the product λ*_tot_*γ, and a time T. Let each local-dynamics function g*^(*i*)^ *be continuously differentiable and bounded (see* subsection 2.3*), and let ε* ∈ (0, 1] *and δ >* 0 *be arbitrary but fixed. For sufficiently large λ*_tot_, *each host’s microbiome abundance vector* **N**^(*i*)^(*t*) *satisfies*

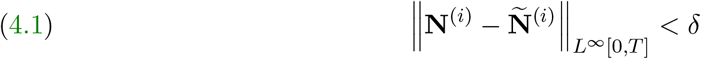

*with probability larger than* 1 − *ε, where*

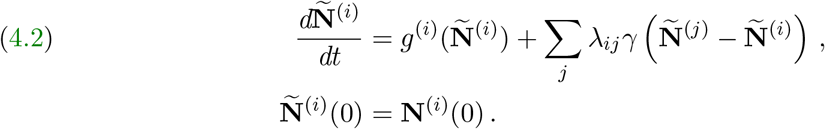

*Proof*. Each local-dynamics function *g*^(*i*)^ is bounded (see subsection 2.3), so there exists a constant *M* such that

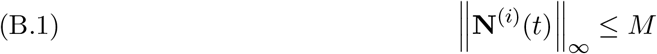

and each entry of **N**^(*i*)^(*t*) is nonnegative for each microbiome abundance vector **N**^(*i*)^(*t*) and all times *t ≥* 0. Each approximate microbiome abundance vector **Ñ** ^(*i*)^(*t*) also satisfies

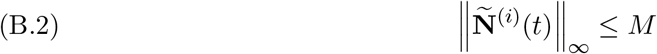

Because

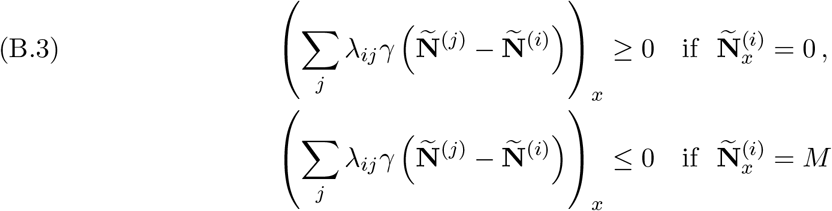

for each entry *x* of **Ñ**^(*i*)^(*t*).

We also assume that each local-dynamics function *g*^(*i*)^ is continuously differentiable and hence continuous. Each host’s microbiome abundance vector **N**^(*i*)^(*t*) is in the region [0, *M*]^*n*^, which is compact, so there exist constants *G* and *F* such that all **N**^(*i*)^(*t*) satisfy the bounds

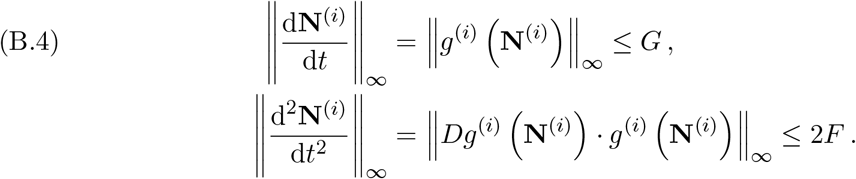

For an interaction involving host *H*^(*i*)^ that occurs at time *t*_*I*_, we choose 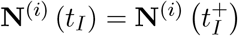. Therefore, **N**^(*i*)^(*t*) is right-continuous at time *t*_*I*_. It is usually not left-continuous at time *t*_*I*_, so it is usually not left-differentiable at time *t*_*I*_.^1^ In such situations, the derivatives that we use in (B.4) are right derivatives. By Taylor’s theorem,

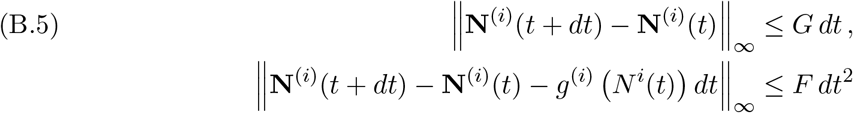

for any interval (*t, t* + *dt*] in which there are no interactions. Each local-dynamics function *g*^(*i*)^ is also Lipschitz continuous. Therefore, there exists a constant *C* such that

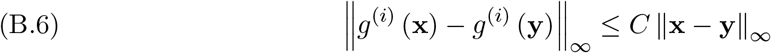

for each *g*^(*i*)^ and all **x, y** ∈ [0, *M*]^*n*^.

Let 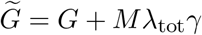, which is a constant because *λ*_tot_*γ* is fixed. For all host microbiome abundance vectors **Ñ** ^(*i*)^(*t*), we have

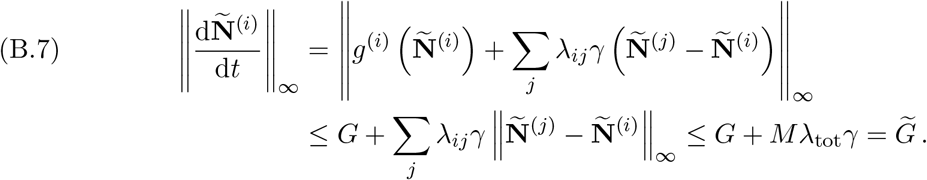

Let 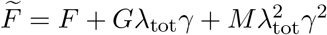. For all **Ñ** ^(*i*)^(*t*), we have

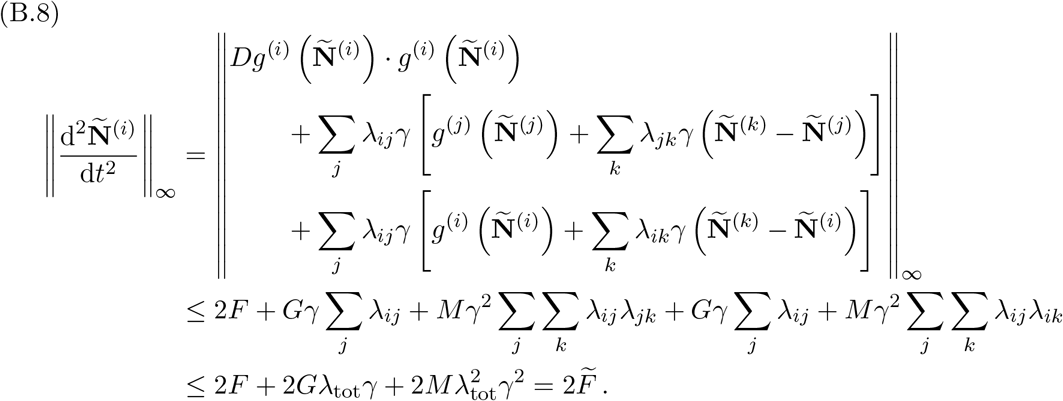

By Taylor’s theorem,

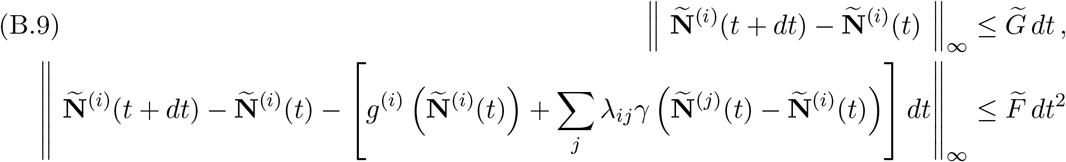

for any times *t* and *t* + *dt*.

We phrased Theorem 4.1 in terms of finding a sufficiently large total-interaction-frequency parameter *λ*_tot_ so that

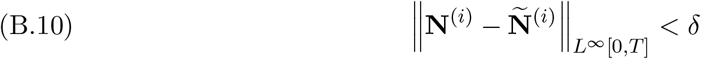

with probability larger than 1 − *ε*. In Theorem 4.1, we assume that the product *λ*_tot_*γ* is fixed, so finding a sufficiently large *λ*_tot_ is equivalent to finding a sufficiently small *γ*. We define the error term

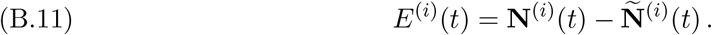

We will show that one can bound each ‖*E*^(*i*)^(*t*) ‖ _*∞*_ with probability larger than 1 − *ε* by a term that involves *γ*. For sufficiently small *γ*, we will show that each

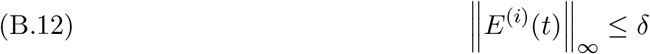

for all *t* ∈ [0, *T*] with probability larger than 1 − *ε*.

Fix a *dt* such that *dt*^4^ *≤ γ ≤* 4 *dt*^4^ *<* 1 and *T/dt* is an integer. This is always possible for sufficiently small *γ*. Let *t*_*k*_ = *k dt*. There are only finitely many *t*_*k*_ in the interval [0, *T*]. The probability that an interaction occurs precisely at any of these *t*_*k*_ is 0. Therefore, for the remainder of this proof, we only consider interactions that occur at times *t* ≠ *t*_*k*_ for any *k*. Under this assumption,

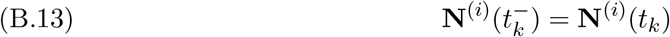

for all *t*_*k*_ [0, *T*].

We now consider how a microbiome abundance vector **N**^(*i*)^(*t*) changes over an interval [*t*_*k*_, *t*_*k*+1_]. Let *L*_*k*_ be the number of interactions that occur in (*t*_*k*_, *t*_*k*+1_). (This interval is open because no interactions occur at any of the *t*_*k*_.) We denote the associated ordered set of interactions by 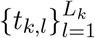. Additionally, we let *t*_*k*,0_ = *t*_*k*_ and *t*_*k,L* +1_ = *t*_*k*+1_, and we define

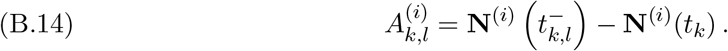

For *l* ∈ {1, …, *L*_*k*_}, let

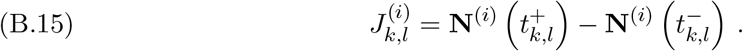

The difference 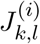 indicates the change in **N**^(*i*)^(*t*) after an interaction at time *t*_*k,l*_. If this interaction does not involve host *H*^(*i*)^, then 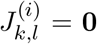. Otherwise, for an interaction between hosts *H*^(*i*)^ and *H*^(*j*)^, we have

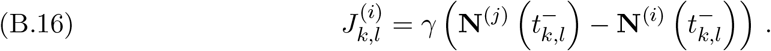

In either case, 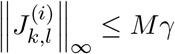. For *l ≥* 2, we decompose the difference 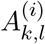 by writing

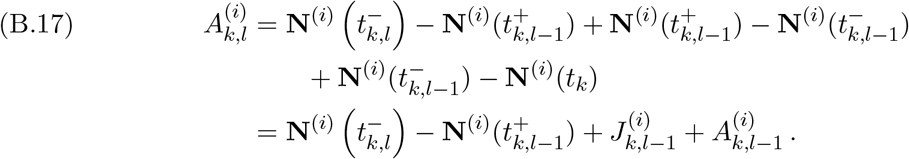

Therefore,

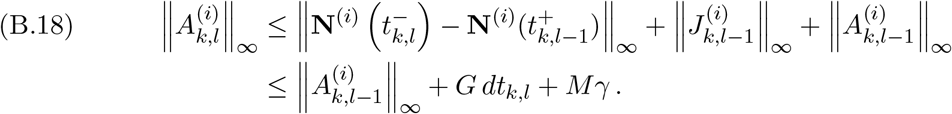

For *l* = 1, we have

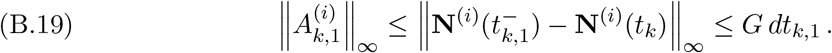

Therefore,

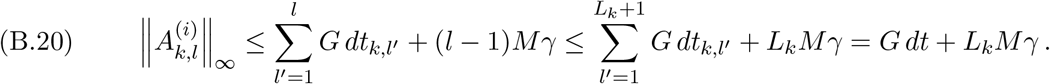

The error between the actual microbiome abundance vector **N**^(*i*)^(*t*_*k*_) and the approximate microbiome abundance vector **Ñ** ^(*i*)^(*t*_*k*_) is

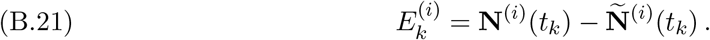

Consider the difference

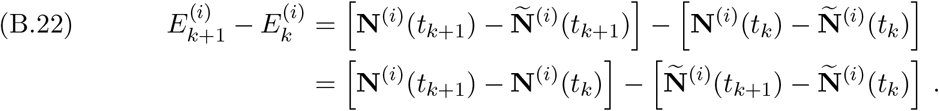

We have

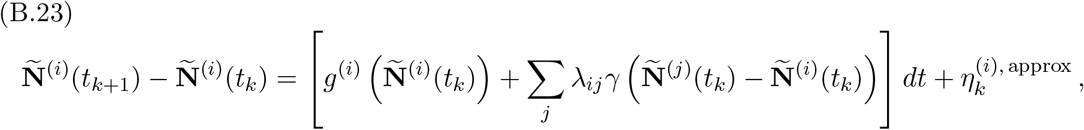

Where

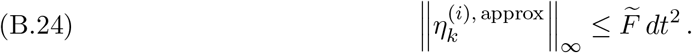

The change of the microbiome abundance vector **N**^(*i*)^(*t*) over the interval [*t*_*k*_, *t*_*k*+1_] is

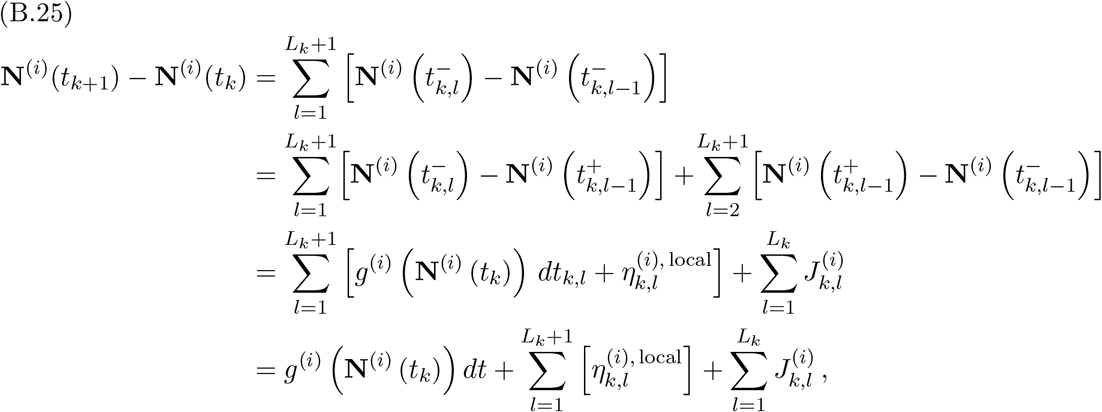

where

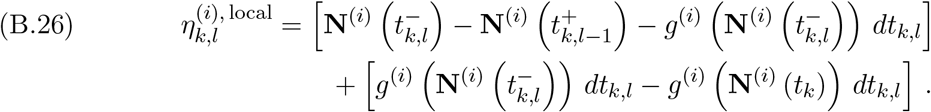

Therefore,

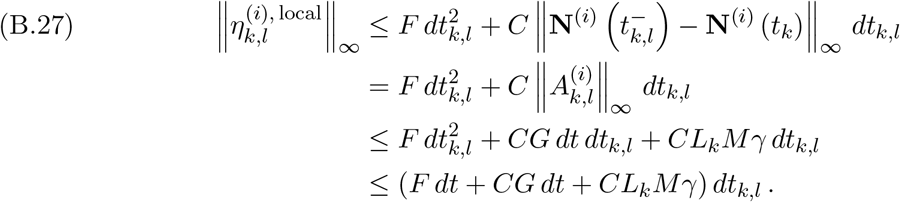

We now consider

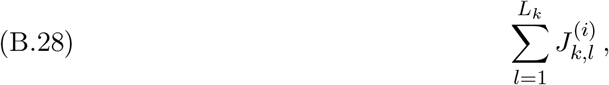

which is the sum of the changes in microbiome abundance vector **N**^(*i*)^(*t*) due to the interactions that host *H*^(*i*)^ has with other hosts. Let

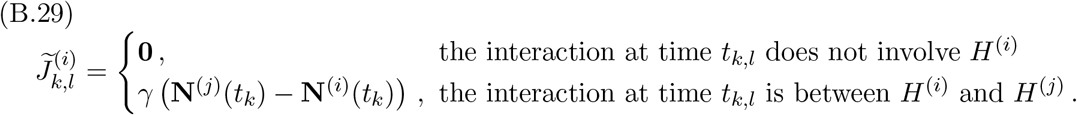

We then have

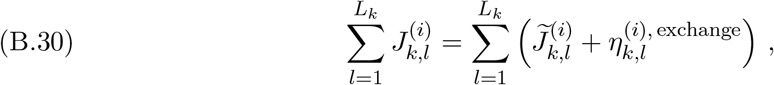

where

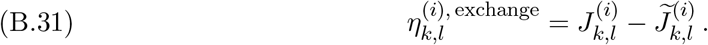

If the interaction at time *t*_*k,l*_ does not involve host *H*^(*i*)^, then 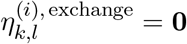. Otherwise, for an interaction at time *t*_*k,l*_ between hosts *H*^(*i*)^ and *H*^(*j*)^, we have

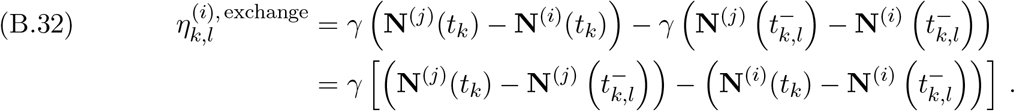

In either case,

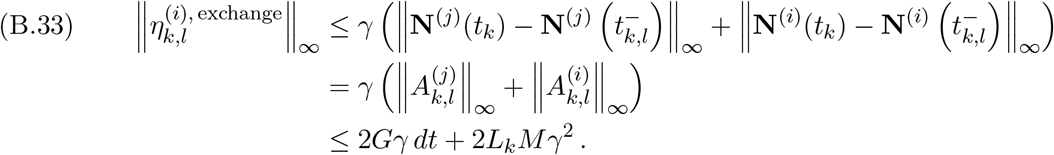

We seek to bound

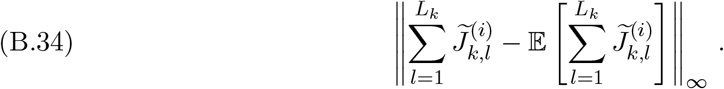

There are two sources of stochasticity in the sum

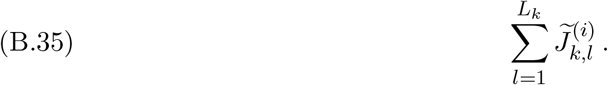

The number *L*_*k*_ of interactions follows a Poisson distribution with mean *λ*_tot_*dt*, and the term 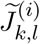 depends on which pair of hosts interacts at time *t*_*k,l*_. Because the sum (B.34) is stochastic, we are only able to bound it with high probability.

The quantity 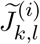 is vector-valued, and we denote an entry *x* of it by 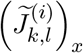. The sum (B.35) is also vector-valued, and we denote an entry *x* of it by 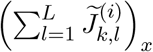. We now calculate the expectation and the variance of each entry of (B.35). The interaction at time *t*_*k,l*_ is between hosts *H*^(*i*)^ and *H*^(*j*)^ with probability *λ*_*ij*_*/λ*_tot_. Therefore,

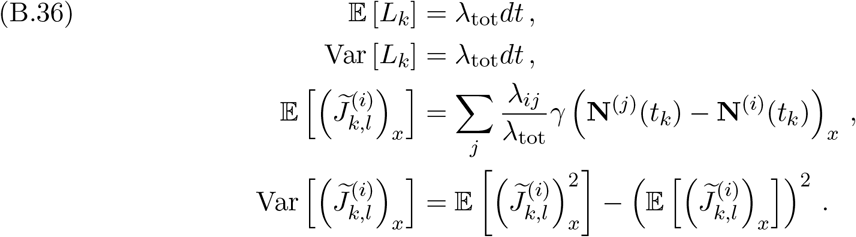

The expectation of each entry of the sum (B.35) is

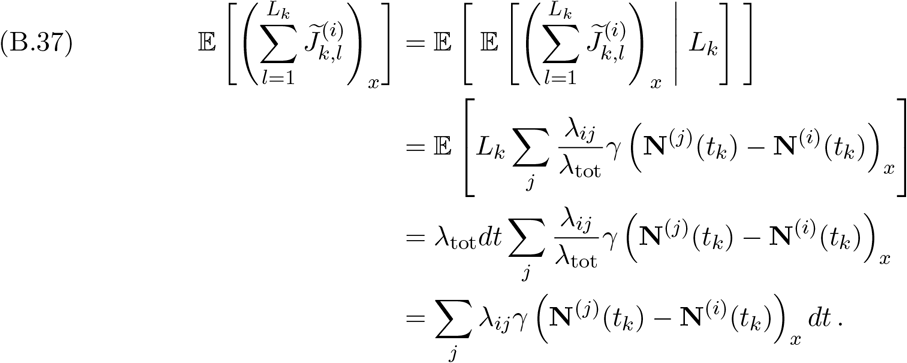

Therefore,

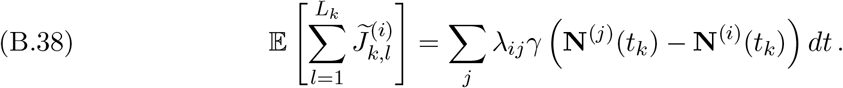

Applying the law of total variance, the variance of each entry of the sum (B.35) is

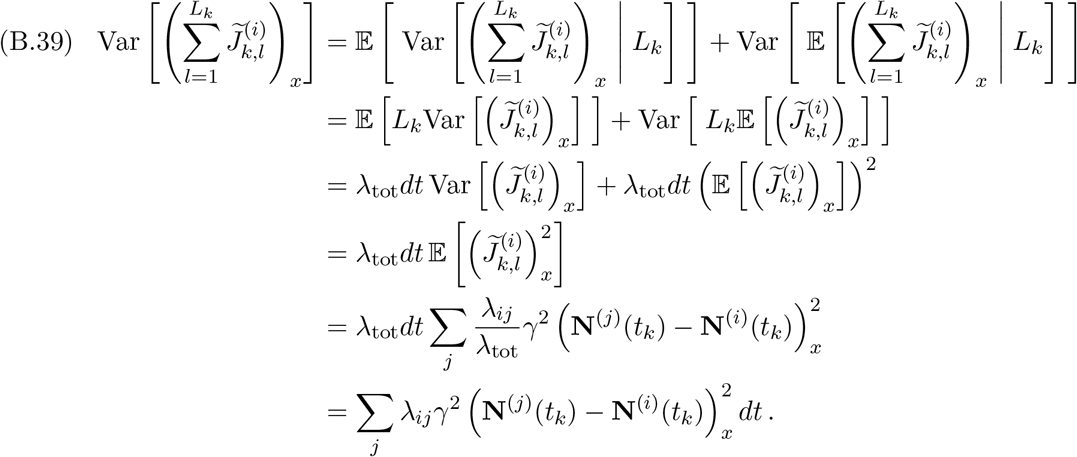

These variances satisfy the bound

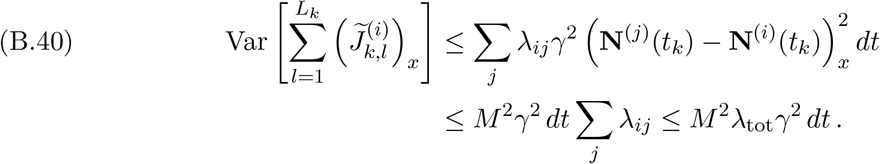

The error due to the stochasticity of the interactions is

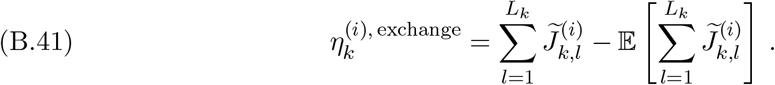

Therefore,

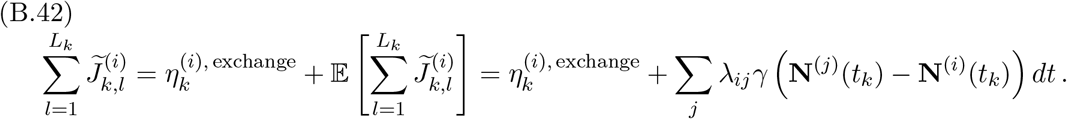

The error (B.41) satisfies

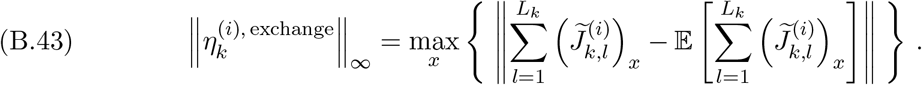

Let *κ* = dim (**N**^(*i*)^) and 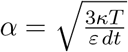. By Chebyshev’s inequality,

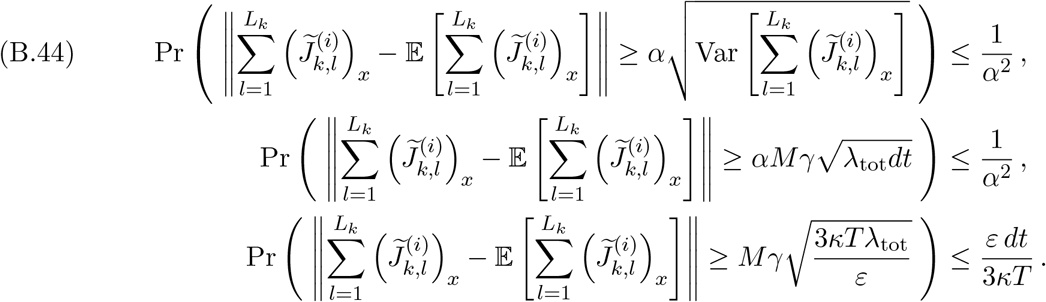

Therefore,

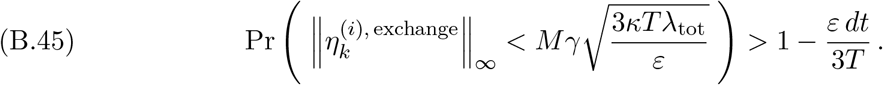

Inserting (B.42) into (B.30) yields

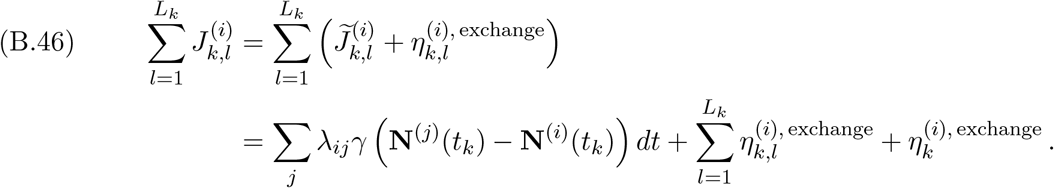

Inserting (B.23), (B.25), and (B.46) into (B.22) yields

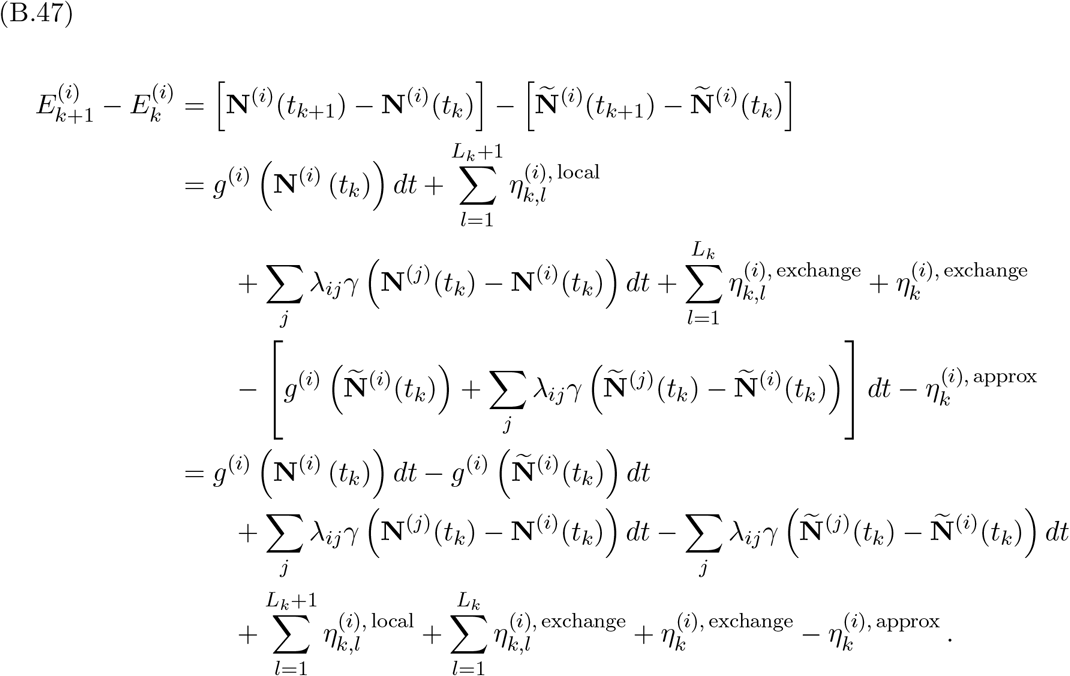

Therefore,

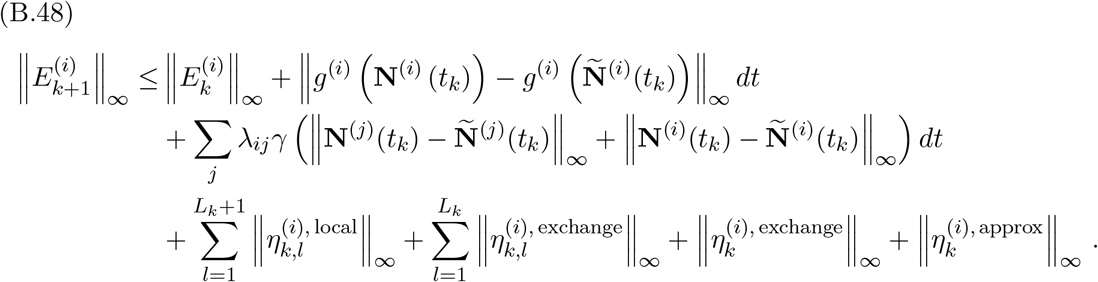

The inequality (B.45) gives a bound for 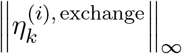 that holds with probability larger than 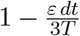. The inequalities (B.27) and (B.33) give bounds for the error terms 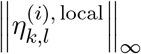 and 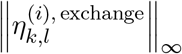, respectively. The latter two bounds depend on the number *L*_*k*_ of interactions in the interval (*t*_*k*_, *t*_*k*+1_). As we discussed above, *L*_*k*_ is stochastic and is distributed as a Poisson random variable with mean *λ*_tot_*dt*. Therefore, we give bounds for *L*_*k*_ that hold with high probability. To obtain these bounds, we first define

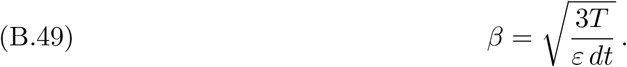

By Chebyshev’s inequality,

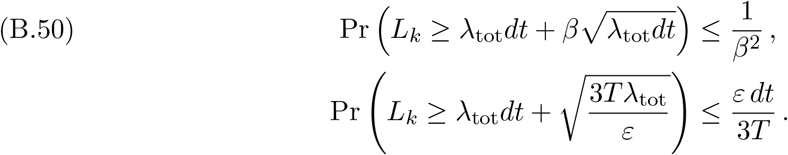

Therefore, with probability at least 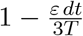, we have the bounds

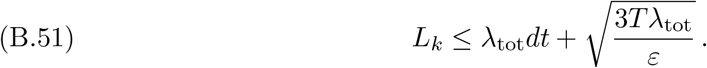

and

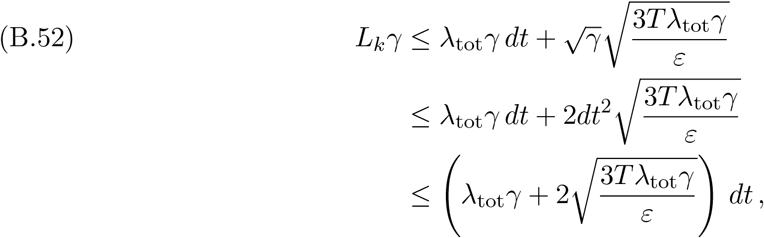

where we use the inequalities *γ <* 4*dt*^4^ and *dt <* 1. With probability at least 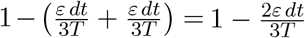, the bounds in (B.45) and (B.52) both hold, yielding

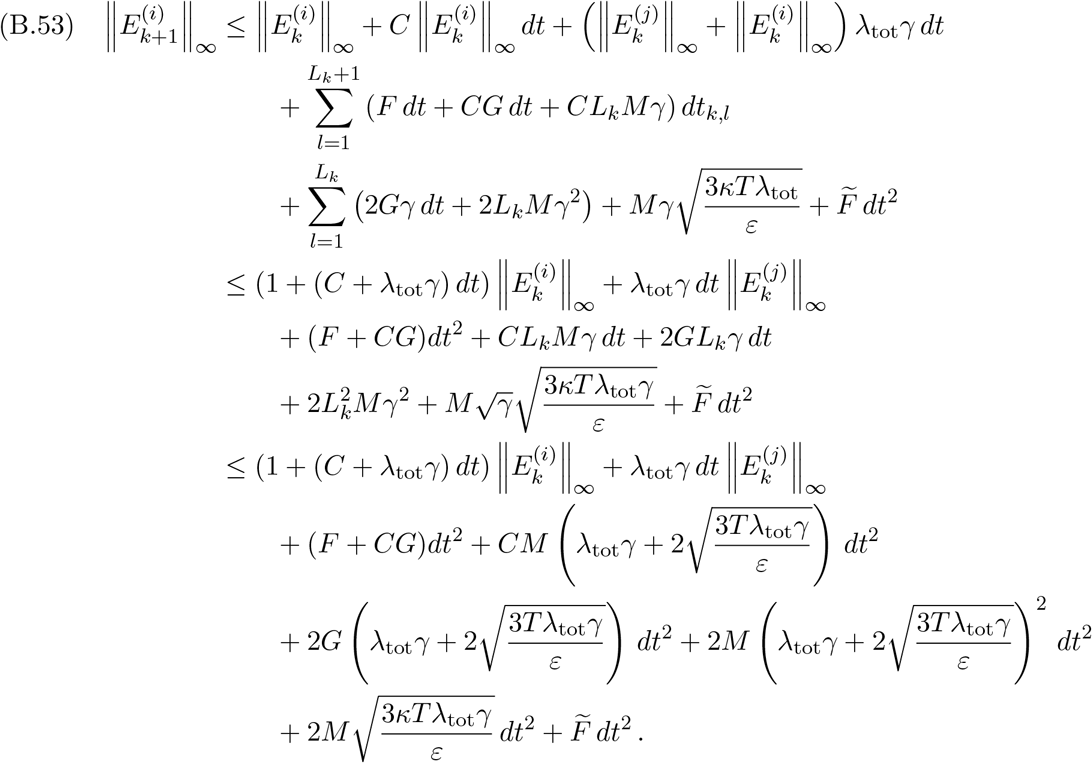

Grouping all of the prefactors of *dt*^2^ into a single constant *Z*, we simplify (B.53) and write

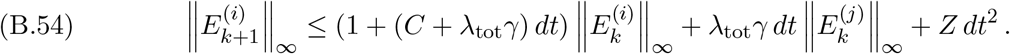

Let

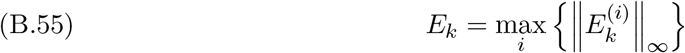

denote the maximum error at time *t*_*k*_. The maximum error at time 0 is *E*_0_ = 0. Let *Y* = (*C* + 2*λ*_tot_*γ*). The maximum error at time *t*_*k*+1_ satisfies the bound

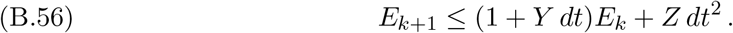

With probability at least 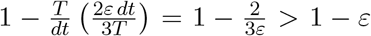, the inequality (B.56) holds for all 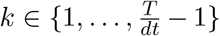. Therefore, with probability larger than 1 − *ε*, the maximum error at time *t*_*k*_ satisfies

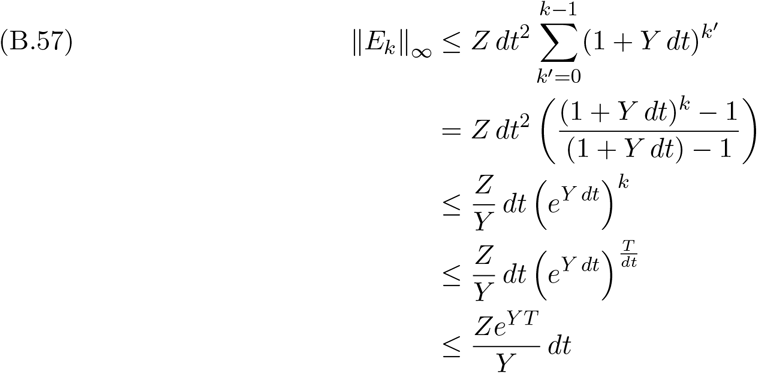

for all 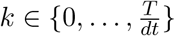

Consider an arbitrary time *t*^*′*^ ∈ (*t*_*k*_, *t*_*k*+1_). For some *l*, we have *t*^*′*^ ∈ [*t*_*k,l*_, *t*_*k,l*+1_). It then follows that

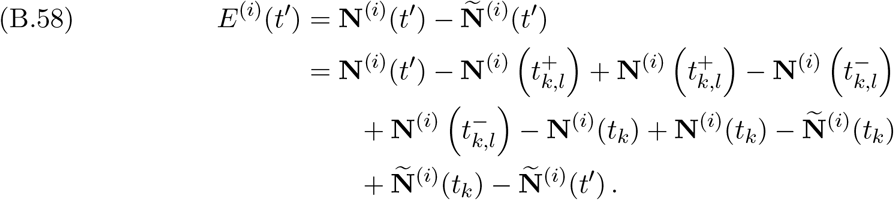

Therefore,

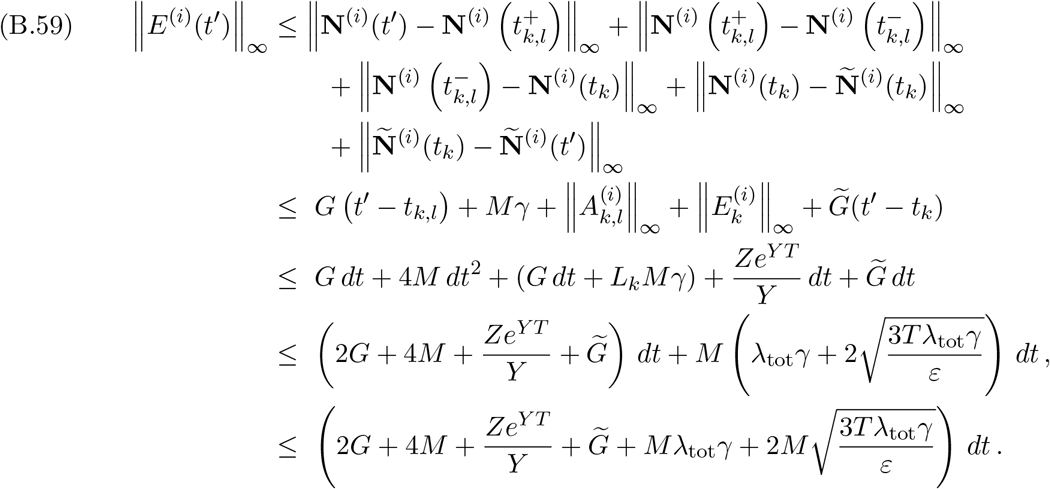

Because 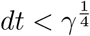, we have

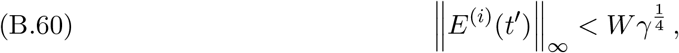

Where

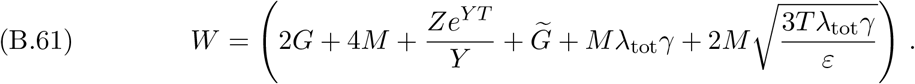

With the inequality

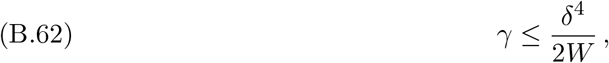

it follows for all *i* and all times *t* ∈ [0, *T*] that

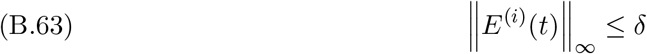

with probability larger than 1 − *ε*.

## Appendix C. Proof of High-Frequency Constant-Strength Approximation Theorem

In this appendix, we prove the HFCSA Theorem (see Theorem 4.2).

### Theorem C.1

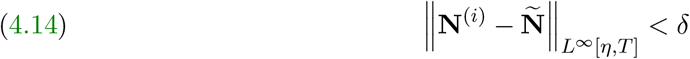

*with probability larger than* 1 − *ε, where*

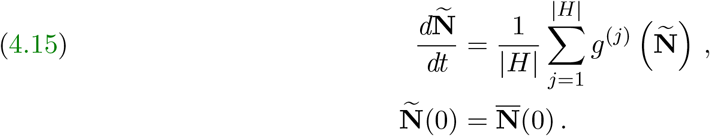

*Proof*. Each local-dynamics function *g*^(*i*)^ is bounded (see subsection 2.3), so there exists a constant *M* such that each entry of **N**^(*i*)^(*t*) is nonnegative and

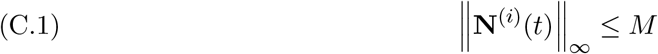

for each microbiome abundance vector **N**^(*i*)^(*t*) and all times *t ≥* 0.

The approximate microbiome abundance vector **Ñ***t*) is the mean of all **N**^(*i*)^(*t*). The dynamics of **Ñ** (*t*) (4.15) is given by the mean of all hosts’ local dynamics. Because each local-dynamics function *g*^(*i*)^ is bounded, so is the mean of each *g*^(*i*)^. Therefore, for all times *t ≥* 0, each entry of **Ñ** (*t*) is nonnegative and

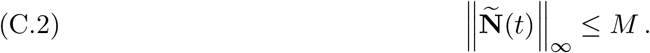

We assume that each local-dynamics function *g*^(*i*)^ is Lipschitz continuous. Therefore, there exists a constant *C* such that

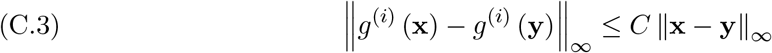

for each *g*^(*i*)^ and all **x, y** ∈ [0, *M*]^*n*^. Because each local-dynamics function *g*^(*i*)^ is continuous (which follows from their Lipschitz continuity) and each microbiome abundance vector **N**^(*i*)^(*t*) is in the compact region [0, *M*]^*n*^, there exists a constant *G* such that

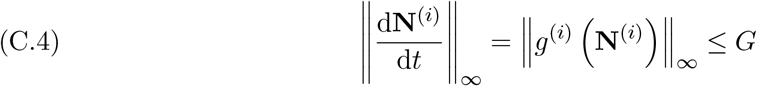

for all **N**^(*i*)^(*t*). For an interaction involving host *H*^(*i*)^ that occurs at time *t*_*I*_, we choose 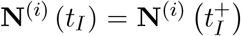. Therefore, **N**^(*i*)^(*t*) is right-continuous at time *t*_*I*_. It is usually not left-continuous at time *t*_*I*_, so it is usually not left-differentiable at time *t*_*I*_.^2^ In such situations, the derivative that we use in (C.4) is a right derivative.

Define the Dirichlet energy

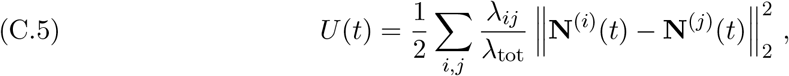

which is nonnegative by construction. Let *ξ >* 0 be arbitrary but fixed. We will show that *U* (*t*) *≤ ξ* with probability larger than 1 − *ε* for sufficiently large *λ*_tot_ and all times *t* ∈ [*η, T*].

Between interactions,

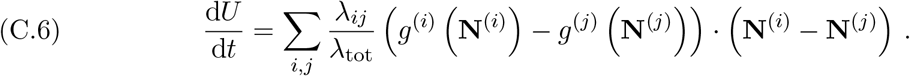

Therefore,

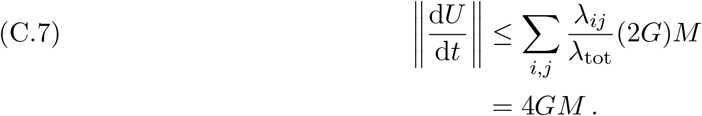

Fix a *dt >* 0 such that *T/dt* is an integer and

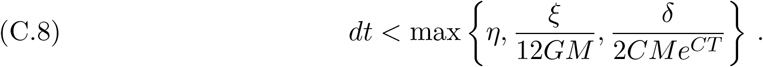

Let *t*_*k*_ = *k dt*. There are only finitely many *t*_*k*_ in the interval [0, *T*]. The probability that an interaction occurs precisely at any of these *t*_*k*_ is 0. Therefore, for the remainder of this proof, we only consider interactions that occur at times *t*≠ *t*_*k*_ for any *k*. Under this assumption,

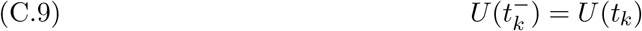

for each *t*_*k*_ ∈ [0, *T*].

We now consider how the Dirichlet energy *U* (*t*) changes over an interval [*t*_*k*_, *t*_*k*+1_]. Let

*L*_*k*_ be the number of interactions that occur in (*t*_*k*_, *t*_*k*+1_). (This interval is open because no interactions occur at any of the *t*_*k*_.) We denote the associated ordered set of interactions by 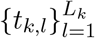. Additionally, we let *t*_*k*,0_ = *t*_*k*_ and 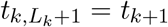, and we define *dt*_*k,l*_ = *t*_*k,l*_ − *t*_*k,l*−1_. For *l* ∈ {1, …, *L*_*k*_}, let

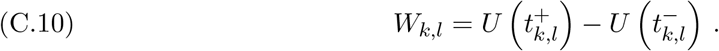

The difference *W*_*k,l*_ is the change in *U* due to an interaction at time *t*_*k,l*_.

We decompose the change in *U* (*t*) over the interval [*t*_*k*_, *t*_*k*_ + 1] by writing

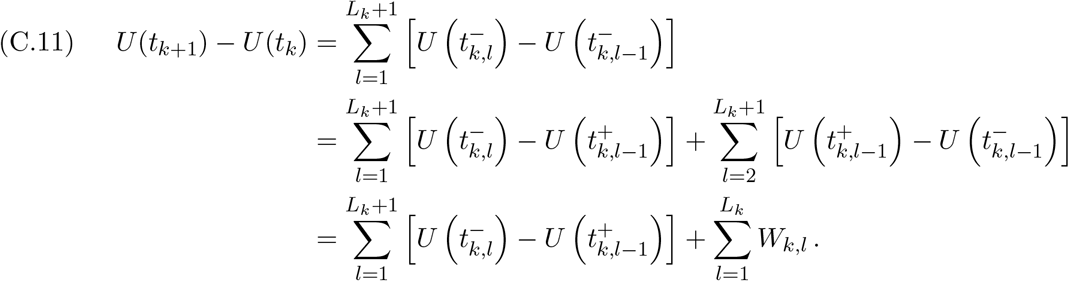

The magnitude of the first sum in (C.11) has the upper bound

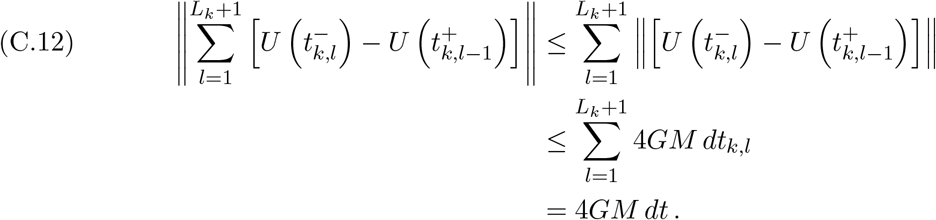

We now consider the sum 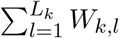. If the interaction at time *t*_*k,l*_ is between hosts *H*^(*i*)^ and *H*^(*j*)^, then

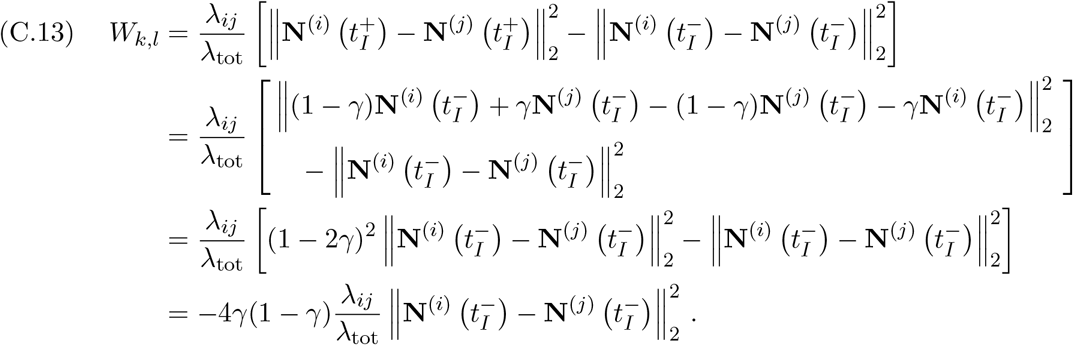

Regardless of which hosts interact, each *W*_*k,l*_ is always nonpositive. We now show by contradiction that the Dirichlet energy *U* (*t*) *≤ ξ/*3 with probability at least 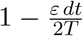 for sufficiently large total-interaction-frequency parameter *λ*_tot_ and some time *t* ∈ [*t*_*k*_, *t*_*k*+1_].

Suppose that *U* (*t*) *> ξ/*3 for all times *t* ∈ [*t*_*k*_, *t*_*k*+1_]. For all *t* ∈ [*t*_*k*_, *t*_*k*+1_], there are then some *i* and *j* such that

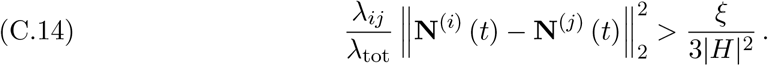

Therefore, at each time 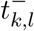 immediately before an interaction, there is some *i* and *j* such that

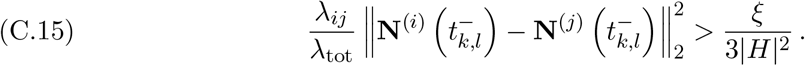

Let

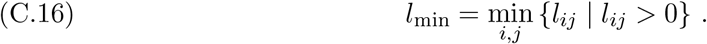

An interaction at time *t*_*k,l*_ occurs between hosts *H*^(*i*)^ and *H*^(*j*)^ with probability *l*_*ij*_ *≥ l*_min_. Therefore, regardless of the previous interactions, each *W*_*k,l*_ satisfies

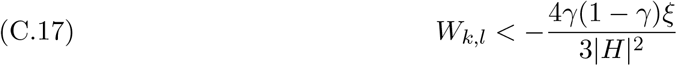

with probability at least *l*_min_.

We define a random variable 𝒲 such that

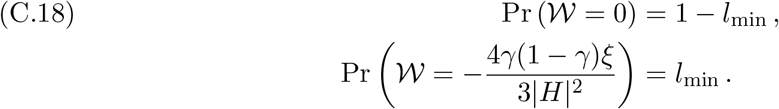

For all *w ≤* 0, we have

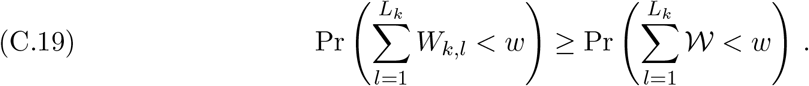

We seek to bound

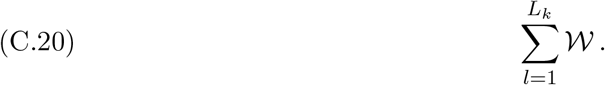

The random variable 𝒲 is stochastic, so we can only find a bound for (C.20) that holds with some probability. The first two moments of 𝒲 are

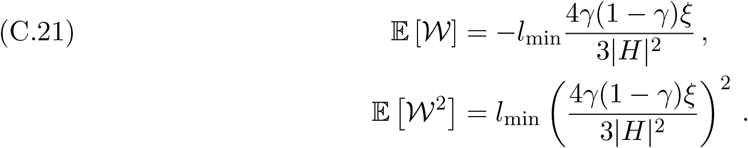

The expectation of the sum (C.20) is

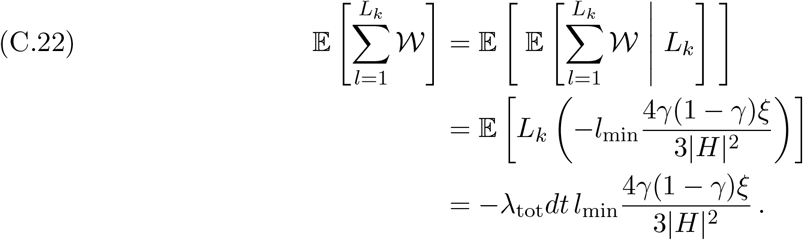

Applying the law of total variance, the variance of the sum (C.20) is

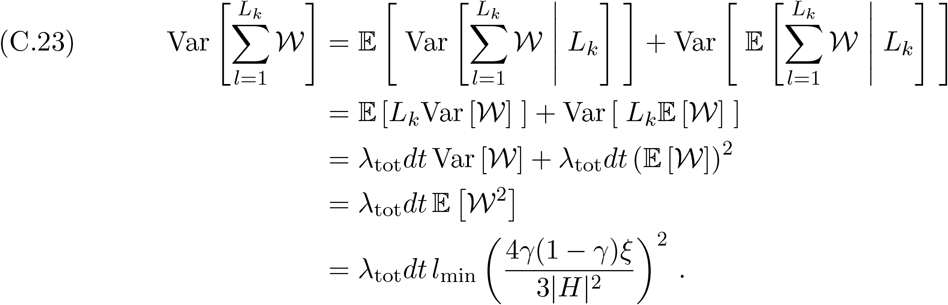

Let 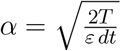. By Chebyshev’s inequality,

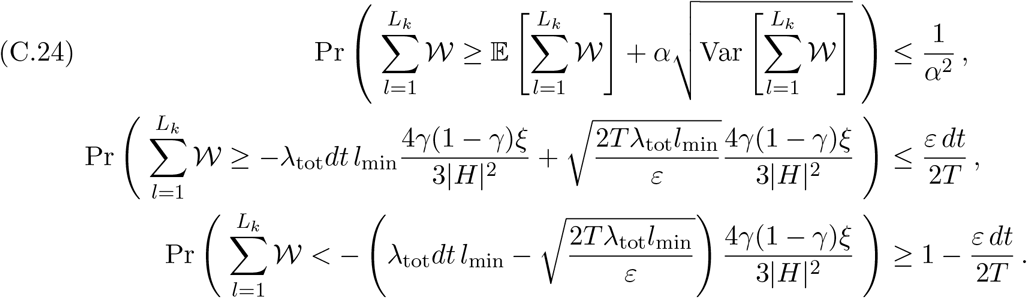

Using the bound (C.19) in (C.24) gives

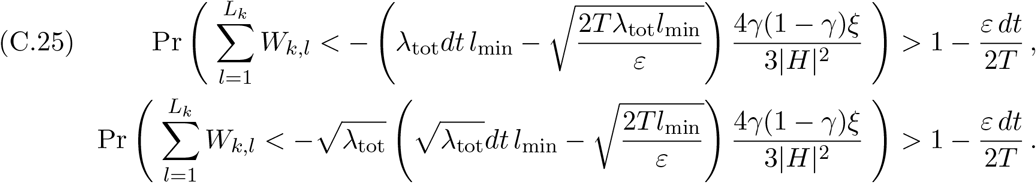

By choosing sufficiently large *λ*_tot_, we can make 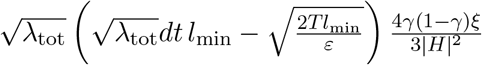 arbitrarily large. In particular, we choose a sufficiently large *λ*_tot_ so that

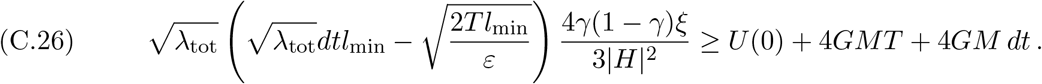

Therefore,

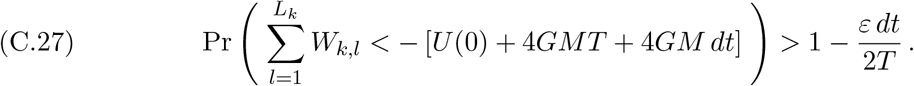

Local dynamics can cause the Dirichlet energy *U* (*t*) to change at a rate of at most 4*GM* per unit time (see (C.7)). Additionally, because each *W*_*k,l*_ is nonpositive, interactions cannot cause *U* to increase. Therefore, for any time *t* ∈ [0, *T*], we have *U* (*t*) *< U* (0) + 4*GMT*. Combining this upper bound, the bound (C.27), and the decomposition of *U* (*t*_*k*+1_) − *U* (*t*_*k*_) in (C.11) yields

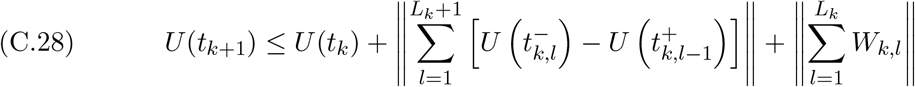

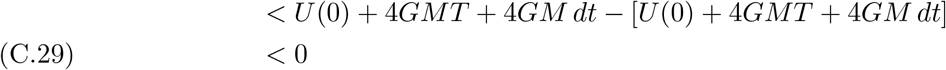

with probability larger than 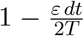. However, *U* (*t*) is always nonnegative. Therefore, with probability larger than 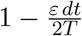, we have a contradiction and there is some time *t*^*′*^ ∈ [*t*_*k*_, *t*_*k*+1_] such that *U* (*t*^*′*^) *≤ ξ/*3. For some *l*^*′*^, we have 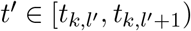. Therefore,

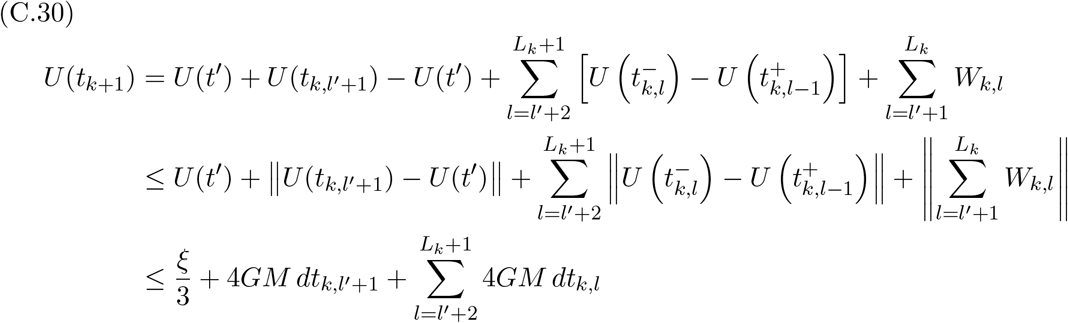

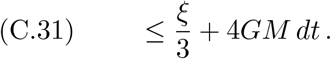

Recall that 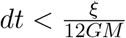 (see (C.8)). We have

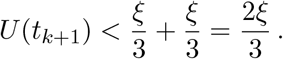

With probability larger than 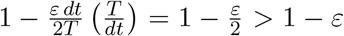*ε*, every *U* (*t*_*k*_) except *U* (0) satisfies *U* (*t*_*k*_) *<* 2*ξ/*3. Consider an arbitrary time *t*^*′*^ ∈ [*t*_*k*_, *t*_*k*+1_], where *k ≥* 1. For some *l*^*′*^, we have 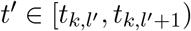. Therefore,

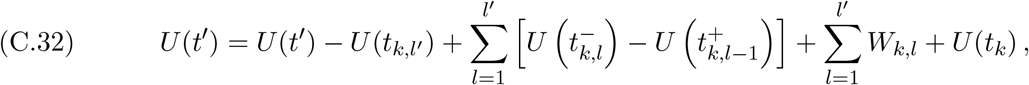

which implies that

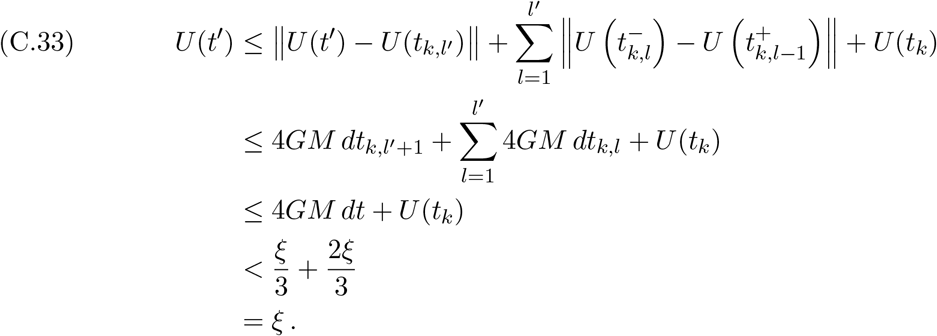

With probability larger than 1 − *ε*, we have *U* (*t*) *< ξ* for all times *t* ∈ [*dt, T*]. Because *ξ* is arbitrary, by choosing a sufficiently large total-interaction-frequency parameter *λ*_tot_, we can make *U* (*t*) arbitrarily small over the interval [*dt, T*] with probability larger than 1 − *ε*. Assume that *U* (*t*) *< ξ* for all *t* ∈ [*dt, T*]. We can then bound the maximum difference between any microbiome abundance vectors **N**^(*i*)^(*t*) and **N**^(*j*)^(*t*). In particular, if *H*^(*i*)^ and *H*^(*j*)^ are adjacent, then

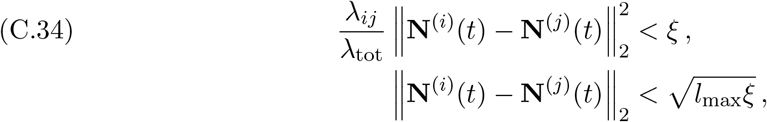

where *l*_max_ = max_*i,j*_ *l*_*ij*_. A shortest path between any two hosts in the interaction network has length at most |*H*| − 1. Therefore, the 2-norm of the difference between each pair of microbiome abundance vectors **N**^(*i*)^(*t*) and **N**^(*j*)^(*t*) satisfies

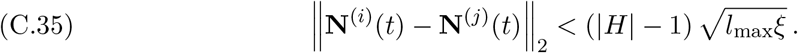

The inequality (C.35) guarantees that the difference between each microbiome abundance vector **N**^(*i*)^(*t*) and the mean microbiome abundance vector 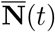 satisfies

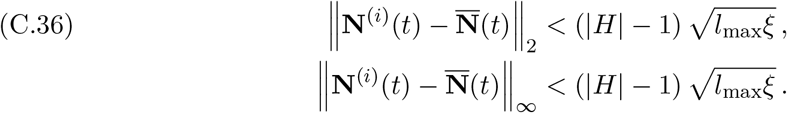

We now bound the magnitude of the difference between the mean microbiome abundance vector 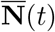 and the approximate microbiome abundance vector **Ñ** (*t*) for all times *t* ∈ [0, *T*]. We refer to 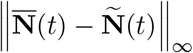 as the *approximation–mean error* at time *t*. At time 0, the approximation–mean error 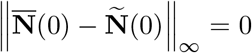 by construction. We bound the approximation–mean error by first bounding its time derivative, which satisfies

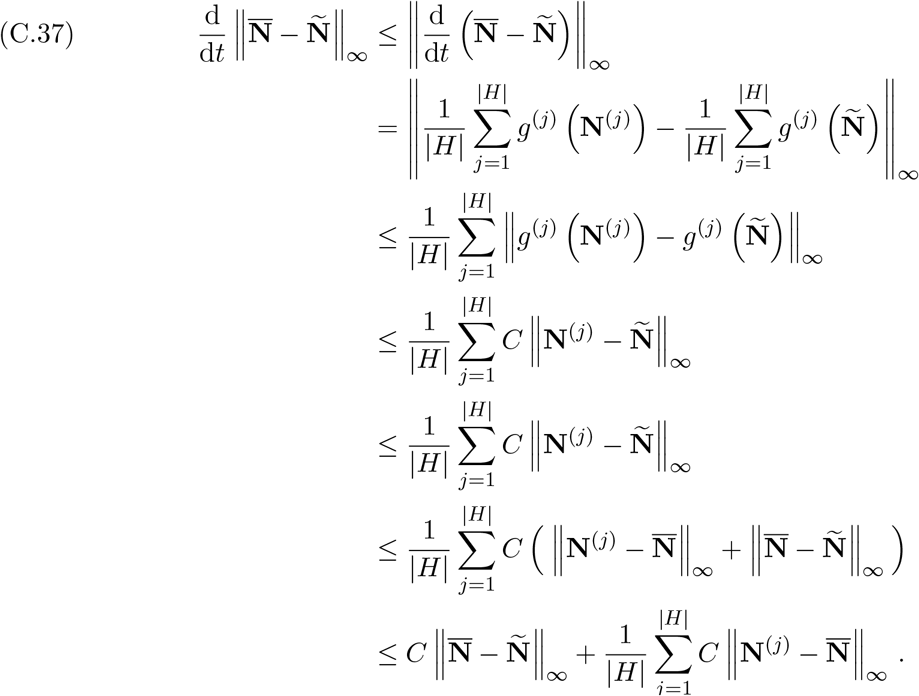

For times *t* ∈ [*dt, T*], each 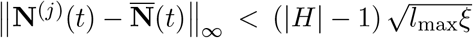. Therefore, on this interval,

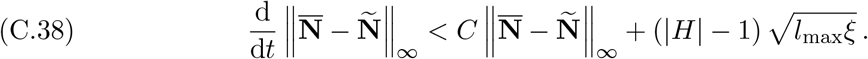

For *t* ∈ [0, *dt*], we obtain a weaker bound for the time derivative of the approximation–mean error:

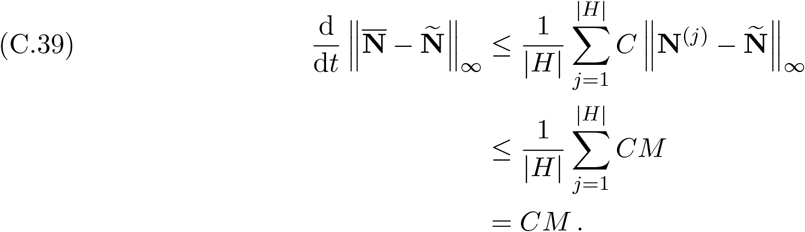

Therefore, for times *t* ∈ [0, *dt*], the approximation–mean error satisfies

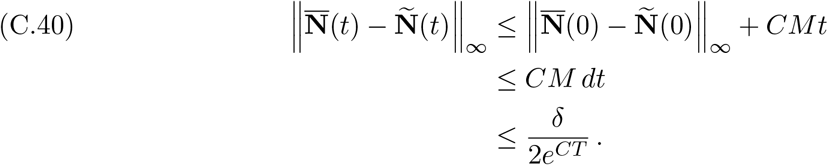

For times *t* ∈ [*dt, T*], we construct a function *u*(*t*) that gives an upper bound of the approximation–mean error. This function *u*(*t*) is the solution of the dynamical system

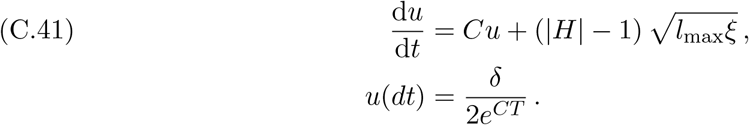

Using an integrating factor to solve for *u*(*t*) yields

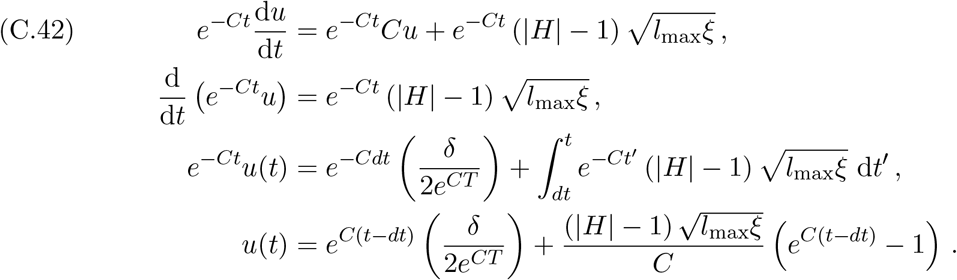

For *t* ∈ [*dt, T*], the function *u*(*t*) satisfies

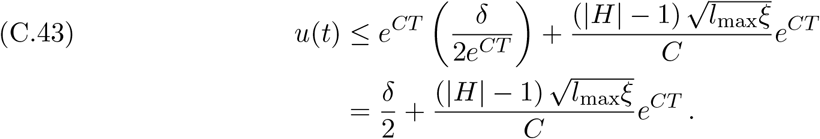

Therefore,

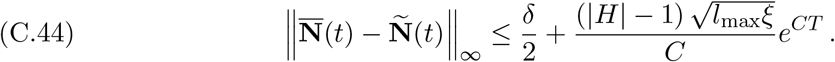

As we discussed previously (see (C.33)), we can make *ξ* arbitrarily small by choosing a sufficiently large *λ*_tot_. Therefore, we make *ξ* sufficiently small so that

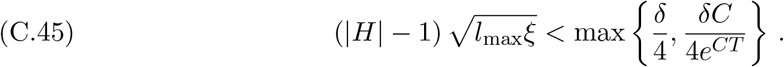

Using the bound (C.45) in (C.44) and (C.36) yields

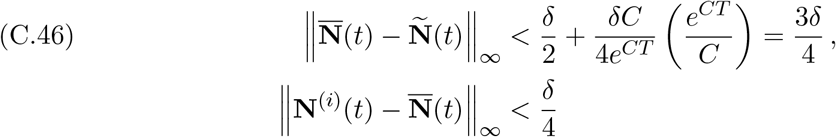

for every *t* ∈ [*dt, T*] with probability larger than 1 − *ε*. Because *dt ≤ η*, for all times *t* ∈ [*η, T*], the difference between each microbiome abundance vector **N** (*t*) and the approximate microbiome abundance vector **Ñ** (*t*) satisfies

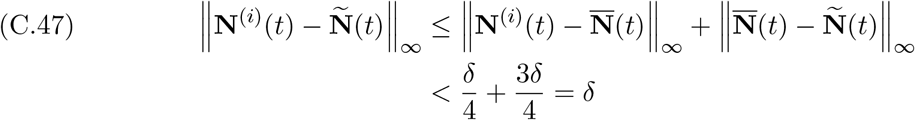

with probability larger than 1 − *ε*.

## Acknowledgements

We thank Kedar Karhadkar, Christopher Klausmeier, Thomas Koffel, Elena Litchman, Sarah Tymochko, and Jonas Wickman for helpful discussions.

## Footnotes

1 The only situation where **N**^(*i*)^(*t*) is left-continuous at time *t*_*I*_ occurs when the host *H*^(*j*)^ with which *H*^(*i*)^ interacts has a microbiome vector 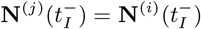.

2 The only situation where **N**^(*i*)^(*t*) is left-continuous at time *t*_*I*_ occurs when the host *H*^(*j*)^ with which *H*^(*i*)^ interacts has a microbiome vector 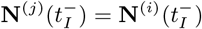.

